# Autopolyploidization affects transcript patterns and gene targeting frequencies in Physcomitrella

**DOI:** 10.1101/2021.06.17.448837

**Authors:** Christine Rempfer, Gertrud Wiedemann, Gabriele Schween, Klaus L. Kerres, Jan M. Lucht, Ralf Horres, Eva L. Decker, Ralf Reski

## Abstract

Qualitative changes in gene expression after an autopolyploidization event, a pure duplication of the whole genome, might be relevant for a different regulation of molecular mechanisms between angiosperms growing in a life cycle with a dominant diploid sporophytic stage and the haploid-dominant bryophytes. Whereas angiosperms repair DNA double strand breaks (DSB) preferentially via non-homologous end joining (NHEJ), in bryophytes homologous recombination (HR) is the main DNA-DSB repair pathway facilitating the precise integration of foreign DNA into the genome via gene targeting (GT). Here, we studied the influence of ploidy on gene expression patterns and GT efficiency in the moss Physcomitrella using haploid plants and autodiploid plants, generated via an artificial duplication of the whole genome. Single cells (protoplasts) were transfected with a GT construct and material from different time-points after transfection was analysed by microarrays and SuperSAGE sequencing. In the SuperSAGE data, we detected 3.7% of the Physcomitrella genes as differentially expressed in response to the whole genome duplication event. Among the differentially expressed genes involved in DNA-DSB repair was an upregulated gene encoding the X-ray repair cross-complementing protein 4 (XRCC4), a key player in NHEJ. Analysing the GT efficiency, we observed that autodiploid plants were significantly GT suppressed (p<0.001) attaining only one third of the expected GT rates. Hence, an alteration of global transcript patterns, including genes related to DNA repair, in autodiploid Physcomitrella plants correlated with a drastic suppression of HR.

## Introduction

The duplication of entire genomes leads to polyploidy and occurs in many cell types and organisms. The resulting polyploid cells and organisms often differ from their progenitors, and are mostly viewed as aberrant or not successful in evolutionary terms. In contrast, recently evidence is accumulating that polyploidization may be an important driving force in evolution as it increases the adaptive potential in stressful conditions (van de Peer et al., 2017), leading to evolutionary innovations and diversification (Walden et al., 2020; Ostendorf et al., 2021).

Sometimes, polyploid cells lose parts of their chromosome set, resulting in aneuploidy. For various eukaryotes including animals, yeast and flowering plants, aneuploidy is in the majority of cases harmful or even lethal (Birchler and Veitia, 2012; Torres et al., 2008). For example, aneuploidy of single cells is regarded as a hallmark of cancer, with about 68% of solid tumours in humans being aneuploid (Duijf et al., 2013; Passerini et al., 2016). It is well established that chromosomal instability causes aneuploidy which drives tumour formation, but there is growing evidence that aneuploidy itself might contribute to tumorigenesis (Ben-David and Amon, 2020). In humans, aneuploidy caused by the addition of one single chromosome, as extensively investigated in the chromosomal-disorder disease trisomy 21, has severe consequences and leads to characteristic phenotypical alterations. Here, the majority of genes on the multiplied chromosome 21 showed a quantitative stoichiometric 1.5 fold increase in expression (Amano et al., 2004). However, in humans with trisomy 21, regions with altered gene expression are not restricted to the affected chromosome, but occur all over the genome, revealing that aneuploidy affects global transcript patterns (Letourneau et al., 2014).

In contrast to aneuploids, euploid organisms deriving from a whole-genome duplication (WGD) are viable and show less phenotypical deviations. The phenotypical effects of WGDs in plants include increased cell sizes provoking a shift in the volume-to-surface ratio of cells as well as increased biomass production (Wu et al., 2012; del Pozo and Ramirez-Parra, 2015). Similar to aneuploidy, a WGD can result in a qualitative change in the expression of selected genes, for example by an upregulation of gene expression stronger than anticipated by the increased gene dosage (Guo et al., 1996), as well as in an unaltered level of gene products, presumably caused by gene dosage compensation mechanisms (Birchler and Veitia, 2012; Shi et al., 2015).

In allopolyploids with their chromosome sets originating from different taxa, a synergy between the chromosome duplication and the hybrid vigor or heterosis effect may occur, associated with increased growth rates, a diverging morphology as well as an improved ability to adapt to new environmental conditions (Comai, 2005; Sattler et al., 2016). Therefore, allopolyploidization is an attractive strategy for the optimization of crop plants in agriculture (Matsuoka, 2011; Behling et al., 2020), and allows them to take over new niches (Cheng et al., 2018). For example, there is molecular evidence for allopolyploidy in some mosses of the genus *Physcomitrium* representing important land pioneers (Beike et al., 2014, Medina et al., 2018). However, in autopolyploids, with chromosome sets from the same taxon, a hybrid vigor effect is lacking and hence the overall impact of a pure WGD on the genome is weaker compared to allopolyploids (Spoelhof et al., 2017). It is unclear to what extent a pure WGD affects gene expression, not only quantitatively due to increased gene dosage but also qualitatively at the global level. A qualitative change in gene expression might contribute to phenotypic effects observed after artificial pure WGDs, like a smaller fruit size in autotetraploid *Hylocereus monacanthus* plants (Cohen et al., 2013) or a reduced viability in stationary phase in isogenic yeast tetraploids (Andalis et al., 2004).

In contrast to animals, land plants undergo an alteration of generations between the haploid gametophyte and the diploid sporophyte. In most cases, this alteration is heteromorphic, i.e. gametophyte and sporophyte are clearly distinguishable in morphology. Whilst the sporophyte dominates in angiosperms, the gametophyte dominates in bryophytes (mosses, liverworts, hornworts). Thus, most bryophytes are haploid in the dominating stage of their life cycle (Reski, 1998a), although diploid or even triploid gametophytes exist, for example in the ecologically important peat mosses (Heck et al., 2021). While the genetic regulator for the developmental switch between haploid and diploid generation has been identified, at least in the moss Physcomitrella (Horst et al., 2016; Horst and Reski, 2016), it remains unclear why these haploid plants are so successful in evolutionary terms, and not prone to excess mutations.

The discovery that Physcomitrella repairs DNA double-strand breaks (DSBs), preferably via the homologous recombination (HR) mechanism (Schaefer and Zrÿd, 1997) may provide an explanation for this enigma. This highly efficient HR machinery facilitates the precise and efficient integration of foreign DNA via gene targeting (GT) with success rates of up to more than 90% in this species (Girke et al., 1998; Kamisugi et al., 2005, 2006; Strepp et al., 1998). Subsequently, highly efficient HR was also described for the moss *Ceratodon purpureus* (Trouiller et al., 2007) and the liverwort *Marchantia polymorpha* (Ishizaki et al., 2013). In contrast, non-homologous end joining (NHEJ) is the preferred mode for the repair of DNA-DSBs in angiosperms. NHEJ relies on a protein complex comprising Ku70, Ku80, DNA-PK_CS_, XRCC4 and DNA ligase 4 (Weterings and Chen, 2008), leads to a random integration pattern of a transgene in the genome, and thereby results in low GT success rates (Britt and May, 2003; Iiizumi et al., 2008). Hence, all attempts to establish efficient GT strategies in seed plants like rice, wheat, maize and tobacco were not particularly successful up to now with reported frequencies as low as 10^-4^ to 10^-5^ (Beetham et al., 1999; Dong et al., 2006; Okuzaki and Toriyama, 2004; Zhu et al., 1999). More recently, the CRISPR/Cas9 system was successfully applied for GT in maize and soybean (reviewed by Steinert et al., 2016), as well as for the realization of agronomic traits in non-browning mushrooms or in maize plants producing high amounts of amylopectin (Qi et al., 2020; Waltz, 2016). However, GT rates are still low and an elaborate screening process is required (Barone et al., 2020; Mao et al., 2019; Schindele et al., 2020).

It is still puzzling why HR is so efficient in some members of the bryophytes. Physcomitrella is a convenient model organism to address this question since it can be easily cultivated under controlled conditions, protocols for precise genetic engineering by GT are well established (Decker et al., 2015) and its genome sequence is available, assembled and annotated (Rensing et al., 2008; Lang et al. 2018). Several possible explanations for the high GT rates in this moss have been discussed, like an altered HR mechanism compared to angiosperms encompassing slight variations in the proteins required for HR or differential expression of their encoding genes (Puchta, 2002; Reski, 1998b; Strotbek et al., 2013). HR-based DNA-DSB repair in Physcomitrella relies on MRE11 and RAD50 (Kamisugi et al., 2012), which are part of a protein complex binding to the ends of broken DNA strands. Targeted knock-out (KO) of the recombinase RAD51 or the SOG1-like protein SOL proved the importance of these proteins in HR and moved DNA-DSB repair to faster but non-sequence conservative repair pathways (Goffová et al., 2019; Markmann-Mulisch et al., 2007; Schaefer et al., 2010). Further, the simultaneous presence of the kinases ATM and ATR, that are also involved in the reprogramming of Physcomitrella leaf cells into stem cells after DNA damage (Gu et al., 2020), are indispensable for GT via HR (Martens et al., 2020). A number of additional proteins have been identified that are favourable but not crucial for GT via HR, like the homology-dependent DSB end-resection protein PpCtIP (Kamisugi et al., 2016) and both subunits of the XPF-ERCC1 endonuclease complex involved in the removal of 3’ non-homologous termini, among others (Guyon-Debast et al., 2019). Additionally, two RecQ helicases possess a crucial distinct function in HR and influence GT frequency, where RecQ6 is an enhancer and RecQ4 a repressor of HR (Wiedemann et al., 2018). Similarly, Polymerase Q (POLQ), containing an N-terminal helicase-like domain, acts as an inhibitor of the HR pathway (Mara et al., 2019).

Hypotheses that are more general were proposed early on: haploidy of the tissue may favour high HR (Schaefer and Zrÿd, 1997), or an unusual cell-cycle arrest may be advantageous (Reski, 1998b). Physcomitrella chloronema cells stay predominantly at the G2/M-boundary (Schween et al., 2003). This cell-cycle phase may be correlated with efficient HR, as HR requires preferentially a sister chromatid as source of the homologous nucleotide sequence that is only available in the late S-phase and the G2-phase of the cell cycle (Heyer et al., 2010; Watanabe et al., 2009). Indeed, B1-type CDKs and B1-type cyclins are regarded to be important regulators of HR in the angiosperm model *Arabidopsis thaliana,* linking the activity of HR to the G2-phase of the cell cycle (Weimer et al., 2016).

A technical way to achieve GT in Physcomitrella is PEG-mediated protoplast transformation. In protoplasts, the recovery from cell-wall removal and isolation of single cells from the tissue is expected to happen in the same period as the integration of the transgene via HR. This is assumed to be completed within the first 72h after isolation before the first cell division of the protoplast (Xiao et al. 2012). Hence, transfected protoplasts are an interesting system to study both of these processes simultaneously. Further, analyses of protoplasts allow insights into plant defence, stress mechanisms and the regeneration of the cell wall (He et al., 2007). In Physcomitrella protoplasts, the primary cell wall was already re-established one day after isolation and after two days, they are partially reprogrammed into stem cells to re-enter the cell cycle. Finally, after three days the majority of the protoplasts have divided and developed into chloronema tissue, which is the basis for the regeneration of the whole plant (Abel et al., 1989; Xiao et al., 2012).

Here, we studied the gene expression patterns in haploid Physcomitrella plants and autodiploid plants with an artificial duplication of the whole genome, created by somatic hybridization via protoplast fusion, and subsequent regeneration of diploid gametophytic plants (Schween et al., 2005b). An analysis of their cell cycle revealed no differences between haploid and autodiploid plants (Schween et al., 2005b). First, we studied the gene expression patterns of haploid regenerating protoplasts in a fine-grained microarray time series using material from different time-points after PEG-mediated protoplast isolation and transfection with a designated GT-construct (Hohe et al., 2004). We observed a gradual regulation of gene expression over time with the highest number of differentially expressed genes (DEGs) at 24h after transfection, many of them related to photosynthesis. In follow-up microarray and SuperSAGE experiments, we compared the gene expression program in haploid and diploid regenerating protoplasts at two selected time-points after transfection with freshly isolated protoplasts and also protonema as controls. The autodiploid lines followed some similar steps of protoplast regeneration as the haploids. However, we identified some time-point-specific differences in gene expression between haploid and diploid protoplasts. Looking for general transcriptomic differences in haploids and autodiploids, we detected 3.7% DEGs in two-factor analyses of the SuperSAGE data in response to the WGD event, though most of the changes were only moderate. Several of these DEGs in regenerating protoplasts after transfection are involved in cell-cycle regulation, DNA-DSB repair and DNA accessibility. Among these DEGs is an upregulated gene encoding the X-ray repair cross-complementing protein 4 (XRCC4), a key player in the NHEJ DNA-DSB repair pathway (Brouwer et al., 2016; Chang et al., 2017; Graham et al., 2016). We subsequently transformed single protoplasts derived from haploid and diploid plants, respectively, with a GT-construct and monitored gene targeting efficiencies in the different plants. As a result, we observed that autodiploid plants attained only approximately one third of the GT rates that would be expected when assuming a similar GT efficiency as in haploid cells. Hence, an alteration of global transcript patterns, especially for genes implicated in DNA repair, in autodiploid Physcomitrella plants correlates with a drastic suppression of homologous recombination.

## Materials and methods

### Plant lines

In this study, a wild type (WT) Physcomitrella (IMSC no. 40001; new species name *Physcomitrium patens* (Hedw.) Mitt., as proposed by Beike et al., 2014 and Medina et al., 2019) was analysed, as well as several lines derived from it. Different haploid and diploid parental lines were derived from WT protoplasts after transformation experiments with a mutagenized cDNA library (Egener et al., 2002; Schween et al., 2005a). For the current study, plants from regenerating protoplasts were selected that had not taken up foreign DNA, as indicated by the absence of the npt II cassette (confirmed by PCR) and did not survive later treatment with antibiotics. While most regenerating plants were haploid, some were polyploid, most likely because of protoplast fusion during the PEG-treatment of the transformation procedure (Egener et al., 2002; Schween et al., 2005a). From this pool of plants, two haploid and three diploid lines were selected: Haploid A, Haploid B as well as Diploid A, Diploid B and Diploid C. Growth on Knop medium differed between the lines but with no significant influence of ploidy (Schween et al., 2005b). Flow cytometric analyses revealed that all Physcomitrella lines analysed here, whether haploid or diploid, remain predominantly at the G2 phase of the cell cycle and only few cells were in the G1 phase (Schween et al. 2005b; Supplemental Figure S1).

We used WT for the construction of a first microarray cDNA library. A second microarray experiment was carried out with WT, Haploid A, Diploid A and Diploid B. A SuperSAGE library was constructed from WT, Haploid A and Diploid A. For quantitative real-time PCR (qRT-PCR), WT, Haploid A and Diploid A were used. We analysed GT rates with Haploid A, Haploid B, Diploid A, Diploid B, and Diploid C.

The characteristics of all moss lines used in this study are compiled in Table 1.

**Table 1.** Characteristics of all Physcomitrella lines used in this study. The cell cycle stage was determined with flow cytometry analysis in Schween et al. (2005b).

| Line | IMSC number | Ploidy | Dominant cell cycle stage | Origin | Experiment |
| --- | --- | --- | --- | --- | --- |
| WT | 40366 (WTIII)<br>40001 (WTIX) | haploid | G2 | wild type | 1<sup>st</sup> microarray,<br>2<sup>nd</sup> microarray,<br>SuperSAGE,<br>qRT-PCR |
| Haploid A | 40369 | haploid | G2 | regenerating transformed protoplasts | 2<sup>nd</sup> microarray,<br>SuperSAGE,<br>qRT-PCR,<br>GT rate |
| Haploid B | 40368 | haploid | G2 | regenerating transformed protoplasts | GT rate |
| Diploid A | 40371 | diploid | G2 | regenerating transformed protoplasts | 2<sup>nd</sup> microarray,<br>SuperSAGE,<br>qRT-PCR,<br>GT rate |
| Diploid B | 40370 | diploid | G2 | regenerating transformed protoplasts | 2<sup>nd</sup> microarray,<br>GT rate |
| Diploid C | 40873 | diploid | G2 | regenerating transformed protoplasts | GT rate |

### Cell culture conditions

For transformation, the different moss lines were cultivated in liquid or on solid modified Knop medium according to Reski and Abel (1985). The material for microarray and SuperSAGE transcriptome studies as well as for qRT-PCR was grown in liquid Knop medium supplemented with microelements (Egener et al., 2002). Cultivation and protoplast isolation were performed as described in Frank et al. (2005).

### Transformation

The GT construct pRKO25.2 (Hohe et al., 2004) contains a 1920 bp cDNA fragment of the cold-responsive gene Pp3c21_180V3, encoding sphingolipid fatty acid desaturase (PpSFD) (Beike et al. 2015; Resemann et al., 2021). This construct contains the coding sequence for neomycin phosphotransferase (npt II) driven by a nopalinsynthase (NOS) promoter and terminator. Transformation, selection and regeneration procedure was performed as described in Frank et al. (2005). Selection was done twice for 2 weeks on medium containing 25 µg/ml G418 (Promega, Mannheim, Germany) starting 2 weeks after transformation with a 2-weeks release period in between. Preparation of material for subsequent RNA isolation and microarray or SuperSAGE analyses was performed with 300,000 protoplasts at different time-points after isolation and transfection with 20 µg of pRKO25.2 construct per time-point.

### PCR analysis

All PCR primers are listed in Supplementary Table T1, a schematic overview of primer locations is given in Supplementary Figure S2. PCR-based analysis of the transgenics was performed according to Schween et al. (2002). The primer combination npt2cdc1-L and npt2cdc1-R was used to detect the npt II cassette. The primers JMLKO25L and JMLKO25R were used to amplify a specific endogenous fragment of 295 bp in WT plants. Homologous integration of the construct into the endogenous genomic locus was monitored using primers JMLKO25-L3 and JMLK2-R5, which were derived from the border of the npt II cassette and the border of the genomic locus at the 5’ end and with primers JMLK2-F3 and JMLKO25-L4, which were derived from the border of the npt II cassette and the border of the genomic locus at the 3’ end, respectively. Two to four independent samples from each plant were tested with all primers to ensure correct identification of KO plants. The significance of ploidy on the transformation results was evaluated with Fisher’s exact test.

### RNA isolation and cDNA synthesis

Total RNA for microarray and SuperSAGE transcriptome studies was isolated from protonema and protoplasts using the RNeasy Plant Mini Kit (Qiagen, Hilden, Germany), applying on-column DNA digestion with DNaseI in accordance with the manufactureŕs protocol. Isolation of total RNA for qRT-PCR was performed in the same manner with protonema as starting material. The DNA digestion was performed as a separate step with DNaseI (ThermoScientific, Darmstadt, Germany) after purification of RNA. cDNA was synthesised using the TaqMan Reverse Transcription Reagents Kit (ThermoScientific) according to the manufacturer’s protocol with oligo(dT) primers. For each of the three technical replicates, cDNA corresponding to 50 ng of total RNA per transcript was used for quantification. A non-transcribed (-RT) control was included to confirm successful DNA digestion. Primers (Supplementary Table T1) were designed with the help of the Roche Life Science Universal Probe Library Assay Design Center (https://lifescience.roche.com). Prior to further analyses, a melting curve analysis was performed for each primer pair. qRT-PCR of protonema samples was conducted using the SensiFastTM SYBR No-ROX Kit (Bioline, Luckenwalde, Germany) in a LightCycler 480 (Roche, Mannheim, Germany). For normalization of variations in cDNA content the reference genes encoding EF1α (Pp3c2_10310V3.1) and TBP (Pp3c12_4720V3.1) were used (Richardt et al., 2010). The relative transcript abundance was calculated in relation to the reference genes with a modified ΔΔC_t_ approach as described in Hellemans et al. (2007).

### Microarray experiments

The microarray experiments were performed with a 90 K whole genome microarray (Combimatrix Corp., Mukilteo, WA, USA) as described previously (Beike et al., 2015; Kamisugi et al., 2016; Wolf et al., 2010). For each time-point per biological replicate, 1.5 µg of RNA were transcribed into cDNA and amplified to aRNA. Subsequently, 5 µg of aRNA were labelled with Cyanin-5 (RNA ampULSe: amplification and labelling kit; Kreatech, Amsterdam, the Netherlands). The resulting labelled aRNA was fragmented (Fragmentation Reagents; Ambion, Austin TX, USA) and hybridised overnight to the microarray following the manufacturer’s instructions. Visualization was performed with a laser scanner (Genepix 4200A; Molecular Devices, Ismaning, Germany) and images were analysed with the Microarray Imager 5.9.3 Software (Combimatrix Corp.). All time-points were analysed in three biological replicates. The microarray slides were stripped with a stripping kit (Combimatrix Corp.) and reused up to four times. The experimental procedure was the same as described previously (Beike et al., 2015; Kamisugi et al., 2016; Wolf et al., 2010).

### Microarray data analysis

Microarray expression values were investigated with the Expressionist Analyst Pro software (v5.0.23, Genedata, Basel, Switzerland). The probe sets were median condensed, and linear array-to-array normalization was applied using median normalization to a reference value of 10,000. Differentially expressed genes were detected using the Bayesian regularised unpaired CyberT test (Baldi and Long, 2001) with Benjamini-Hochberg false discovery rate correction and a minimum |log_2_ fold change| > 1 (Richardt et al., 2010). A false discovery rate of q < 0.05 was taken as cut-off for the first microarray time series experiment. For the second microarray time series experiment p < 0.001 was chosen for the comparison of gene expression between the ploidy levels and for the comparison of gene expression between different time-points in regenerating protoplasts. K-means clustering with k = 2 identified upregulated and downregulated genes. An overview of the plant lines and sample sources used for the different comparisons to compute DEGs is compiled in Supplementary Table T2.

### SuperSAGE library construction

SuperSAGE libraries were constructed by GenXPro (Frankfurt am Main, Germany) following a protocol based on Matsumura et al. (2010) as described by El Kelish et al. (2014) with the implementation of GenXPro-specific technology and improved procedures for quality control as well as specific bias proved adapters for elimination of PCR artefacts (True-Quant methodology). In total, 17 SuperSAGE libraries (including replicates) were constructed from 11 biological samples. The biological samples encompass: The transcriptome of Haploid A and Diploid A after protoplast isolation (0h) and 4h and 24h after transfection; haploid as well as diploid protonema mRNA in duplicates; transcript data of WT protoplast from 0h, 4h and 24h with triplicates for 4h and 24h. A detailed overview of the libraries is provided in Supplementary Table T3.

### SuperSAGE data analysis

The quality of the processed libraries was checked with FastQC (v0.11.4, Andrews, 2010) and reads were mapped with HISAT2 (v2.0.3, Kim et al., 2015) to the V3 assembly of the *P. patens* genome (Lang et al., 2018) in the Galaxy platform (Freiburg Galaxy instance, http://galaxy.uni-freiburg.de, Afgan et al., 2016). Mapping parameters allowed for no mismatches and only known splice sites were considered. A count table was constructed from the mapped reads using the featureCounts (v1.4.6.p5, Liao et al., 2014) tool from the Galaxy platform by counting all the reads mapped to exons or untranslated regions of each gene. Multiple alignments of reads were allowed, while reads with overlaps on the meta-feature (gene) level were disregarded for the construction of the count table. For specific parameters, see Supplementary Table T4 and Supplementary Table T5. Statistical analysis for differential gene expression was performed by pairwise comparison of library count tables using GFOLD (v1.1.4, Feng et al., 2012) and by two two-factor analyses with the DESeq2 package in Galaxy with default parameters (Galaxy Version 2.11.40.6, Love et al., 2014). In the two-factor analyses, ploidy-dependent gene expression was determined in the presence of tissue as secondary factor. All libraries originating from protonema and different protoplast material were used as input for the first two-factor analysis and the libraries of mock transformed WT protoplasts at 4h and 24h were considered as replicates to the libraries of transformed WT protoplasts at the corresponding time-points. Only libraries derived from protoplasts of the lines WT and Diploid A were considered for the second two-factor analysis. In GFOLD analysis, genes with a GFOLD(0.01) value (representing the log2 fold change of gene expression adapted for adjusted p-value, Feng et al., 2012) of < −1 or > 1 were considered to be differentially expressed whereas in DESeq2 analysis genes with a |log2 fold change| > 1 and an adjusted p value < 0.1 were considered as differentially expressed. Further data exploration was performed using functions from SAMtools (v1.3.1, Li et al., 2009).

### Computational analysis of DEGs

Annotation of DEGs was obtained using Phytozome (v12.1.5, Goodstein et al., 2012) and the PpGML DB (Fernandez-Pozo et al., 2020). For the computation of the overlap between DEGs identified in the microarray and SuperSAGE data, and to generate a combined set of DEGs comprising all DEGs from both technologies, gene IDs of DEGs identified in the second microarray experiment were converted to Physcomitrella V3.3 IDs (Lang et al., 2018). If one ID mapped to several genes of the V3.3 annotation all of them were considered as DEGs. In case the IDs of multiple DEGs mapped to the same V3.3 ID the mean of the log2 fold change values was taken. Similarly, in the comparison between the DEGs identified in our study and DEGs found by Xiao et al. (2012), Physcomitrella V1.6 IDs were translated into Physcomitrella V3.3 IDs. In both cases, genes with no correspondence in the V3.3 annotation were neglected. This amounted to a maximum of 50 out of 2245 DEGs that were not analysed further. Only genes that are contained in the main V3 genome according to the annotation file downloaded from PpGML DB were included in the lists of DEGs presented here. Expression data of specific genes in different developmental stages of Physcomitrella were obtained from the PEATmoss website (Fernandez-Pozo et al., 2020). Genes that are relevant for DNA-DSB repair or for DNA repair in general were identified using biological process (PB) GO terms from the current V3.3 annotation obtained from PpGML DB and an in-house list with repair relevant genes. The principle component analysis (PCA) of SuperSAGE libraries, the gene ontology (GO) enrichment analysis and the visualization of the word cloud with enriched GO terms were carried out in R (v3.6.3, R core team, 2020). For the PCA the R package DESeq2 (v1.24.0; Love et al., 2014) was used, the word cloud with enriched GO terms was created with the R package tagcloud (v0.6, Weiner, 2015) and the word size scales with the negative log2 of the adjusted p value. The GO enrichment analyses were performed with the R package clusterProfiler (v3.12.0, Yu et al., 2012) using a p value cut-off of 0.005 and a q value cut-off of 0.2. The minimal size of genes annotated by ontology term for testing (minGSize) was set to 1 and the maximal size of genes annotated for testing (maxGSize) was set to 1000. As universe for microarray data, all genes on the microarray were taken, whereas the universe for data from SuperSAGE contained all genes of the main Physcomitrella V3 genome. Redundant GO terms were removed afterwards using the *simplify* method from clusterProfiler with default parameters. Generation of several figures and processing of tables with DEGs was performed in Python3 (v3.8.5, Van Rossum and Drake, 2009) using the packages Matplotlib (v3.2.1, Hunter, 2007), NumPy (v1.18.4, Harris et al., 2020), pandas (v1.0.3, McKinney, 2010; The pandas development team, 2020), rpy2 (v3.3.3, Gautier, 2010), seaborn (v0.10.1, Waskom et al., 2020), and pyvenn (https://github.com/tctianchi/pyvenn).

## Results

### Regenerating protoplasts exhibit a time-dependent gene expression pattern

We generated a transcriptomic time series using microarray technology to investigate how gene expression is adjusted during the regeneration of transfected Physcomitrella protoplasts, and to identify those time-points during protoplast transformation with the strongest alterations in gene expression. We assume that transformation of the genome (= integration of the heterologous DNA) is completed before the first cell division of the protoplast, that happens under our conditions within the first 72h after isolation. The 90K whole genome microarray used in this study represents all identified or predicted Physcomitrella gene models of the genome assembly V1.2 (Rensing et al., 2008). Data were generated for WT samples at six time-points: freshly isolated (0h) protoplasts as well as for protoplasts 1h, 4h, 6h, 24h and 72h after transfection.

A pairwise comparison of each time-point after transfection to freshly isolated protoplasts (0h) revealed two maxima of differentially expressed genes (DEGs) at 4h (207 DEGs) and 24h (1064 DEGs, Fig. 1a), whereas gene expression appeared to be unaltered shortly after transfection in 1h old protoplasts. Most of the DEGs at 6h were also differentially expressed at 4h (Fig. 1b). These include a gene encoding glyceraldehyde-3-phosphate dehydrogenase (Pp3c21_9380), a key player in glycolysis, and the genes encoding malate synthase (Pp3c20_22510) and isocitrate lyase (Pp3c7_2470) respectively, which are enzymes of the glyoxylate cycle (Supplementary Excel sheet1). 24h after transfection, 933 additional DEGs that were not identified at any of the earlier time-points were detectable. 10 of the DEGs detected at 4h and 6h were not significantly differentially expressed at 24h anymore. Nearly half of the DEGs found at 72h were specific for this time-point. However, 9 genes that were exclusively differentially expressed at the beginning of protoplast isolation at 4h but not 6h and 24h were now again among the DEGs, for example the genes encoding 4-coumarate-CoA ligase (Pp3c19_13170) and phenylalanine ammonia lyase (Pp3c2_30610). 18 out of the total 1385 DEGs were constantly differentially expressed at each time-point between 4h and 72h, three of them being associated with the plant hormone gibberellin that regulates developmental processes, also in Physcomitrella (Vandenbussche et al., 2007). These are Pp3c5_4920 and Pp3c4_2230, both encoding 2-oxoglutarate and Fe(II)-dependent oxygenases involved among others in gibberellin biosynthesis, as well as Pp3c1_15680 that encodes a homologue to the *A. thaliana* GRAS family protein RGL1. We also detected and jasmonic acids, at up to three time-points. Another gene with a constant differential regulation over time was Pp3c19_6540, which encodes a catalase that acts against oxidative stress by degradation of H_2_O_2_. Several other stress-related genes were differentially expressed at one specific or multiple time-points during protoplast regeneration.

**Fig. 1.**
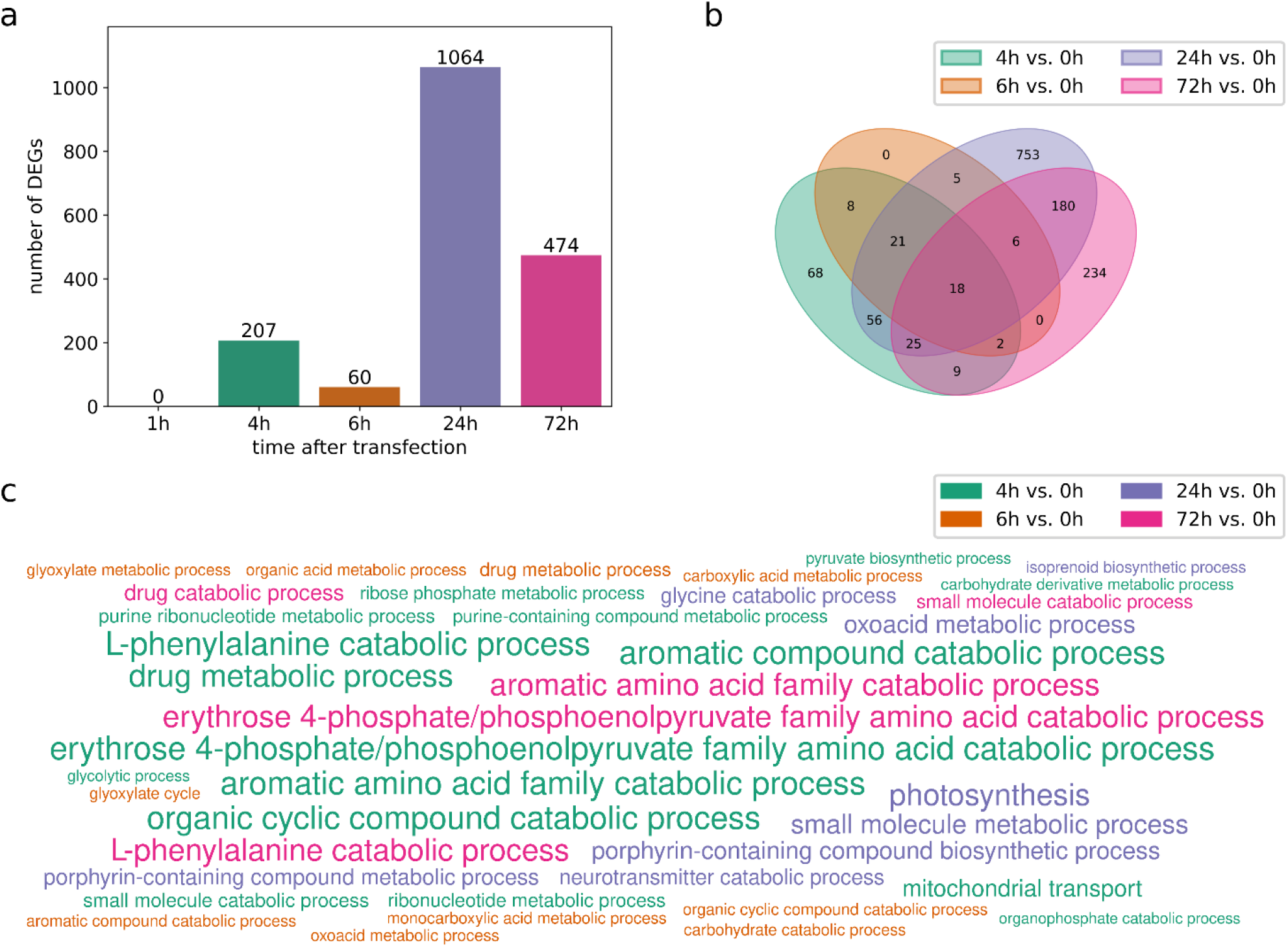
Time series of DEGs in Physcomitrella WT protoplasts 1h, 4h, 6h, 24h and 72h after transfection compared to freshly isolated (0h) protoplasts based on microarray data. DEGs are filtered for q < 0.05. (a) Number of DEGs at each time-point. Two maxima of DEGs are apparent after 4h and 24h, respectively. (b) Overlap of DEGs from each time-point. (c) Significantly enriched biological process GO terms (p < 0.001, q < 0.2). The word size scales with the negative log2 of the adjusted p value.

To gain a deeper insight into protoplast regeneration, we performed a gene ontology (GO) enrichment analysis for those time-points where DEGs were identified (Fig. 1c). We considered terms with p < 0.001 and q < 0.2 as significantly enriched and reduced the number of resulting GO terms by selecting a representative term amongst similar terms. A strong enrichment of genes associated with erythrose 4-phosphate/phosphoenolpyruvate family amino acid catabolic process, L-phenylalanine catabolic processes and aromatic amino acid family catabolic processes was observed at 4h and at 72h (Supplementary Excel sheet 2). Only at 4h, genes with the GO term glycolytic process were enriched, while the GO terms drug metabolic process, organic cyclic compound catabolic process and aromatic compound catabolic process that were strongly enriched at 4h were also enriched at 6h, however to a lower extent. Other enriched GO terms at 6h were for example carbohydrate catabolic processes and glyoxylate cycle. At 24h, mostly the expression of genes with the GO-term photosynthesis altered. Other strongly enriched GO terms in the DEGs at 24h were porphyrin-containing compound biosynthetic process, oxoacid metabolic process and small molecule metabolic process. Further, we found an enrichment of ammonia-lyase activity and ammonia-ligase activity both at 4h and 72h, while we observed an enrichment of aminomethyltransferase activity at 24h.

### A high number of DEGs in haploid versus diploid protoplasts at 24h

To investigate if haploid and diploid protoplasts behave differently during regeneration, we generated transcriptomic data of two haploid and two diploid lines. We analysed protonema, untransformed protoplasts (0h), and protoplasts at 4h and 24h after transfection (Fig. 2, Supplementary Table T6). For a more detailed follow-up analysis, we additionally applied the SuperSAGE sequencing technology. Compared to microarray platforms, SuperSAGE has the advantage that sampling is based on sequencing rather than hybridization of RNA and as a consequence, sequences do not need to be known *a priori*. Furthermore, sequenced reads can be directly mapped to the genome and gene expression is quantified by direct counts of transcript abundance, thus eliminating background noise that exists in microarrays, leading to increased sensitivity and improved gene expression quantification. Altogether, we generated 17 SuperSAGE libraries (Supplementary Table T3) from untransformed protoplasts (0h), and protoplasts at 4h and 24h after transfection as well as from protonema tissue of two haploid lines (WT, Haploid A) and one diploid line (Diploid A). 4 of the libraries (2 WT samples at 4h and 24h respectively) were derived from mock transformants subjected to the whole transfection procedure but using water instead of the GT construct. Pairwise comparison between SuperSAGE libraries were performed with GFOLD, an algorithm especially developed for approaches when only few replicates are available.

**Fig. 2.**
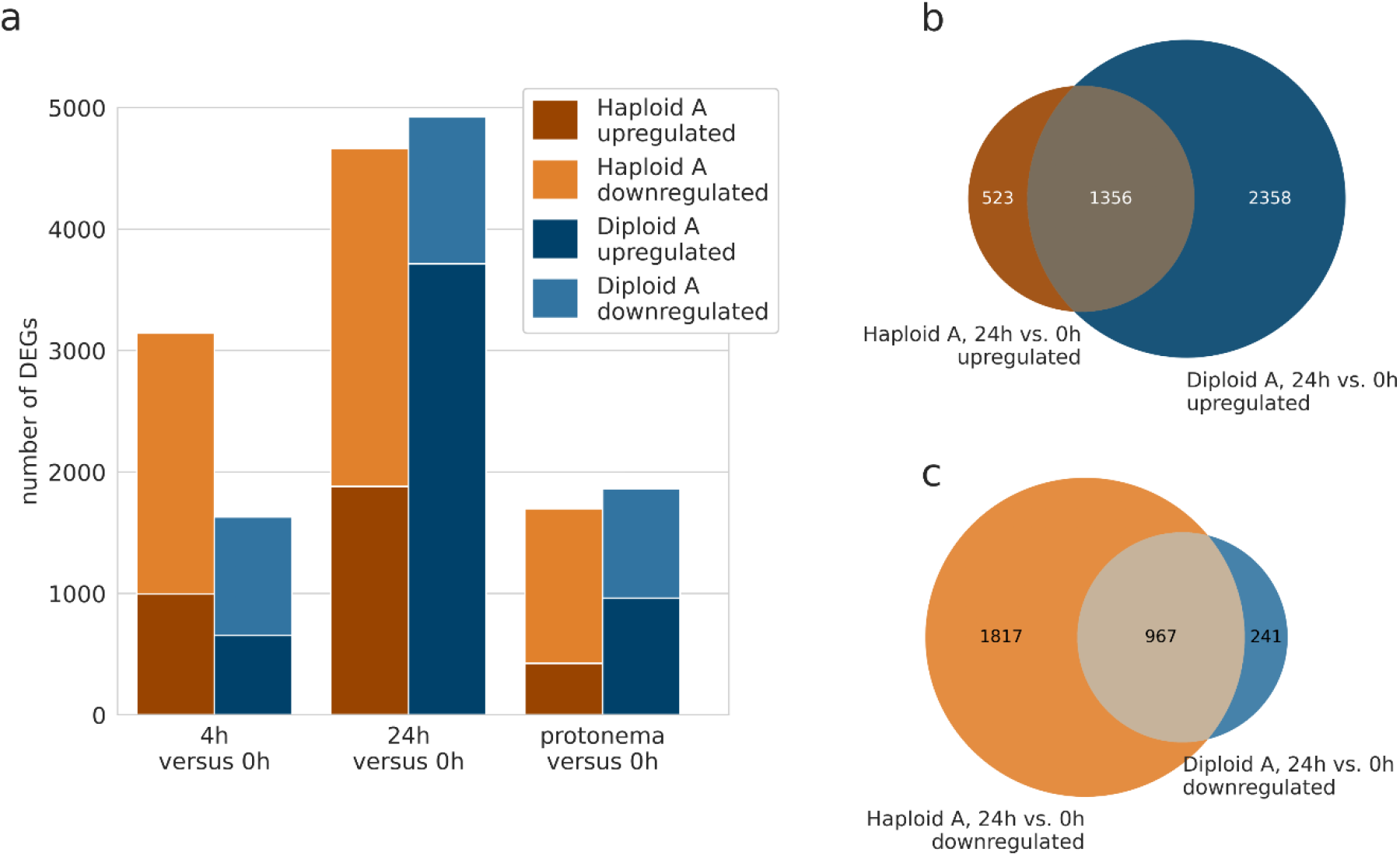
Number of genes being upregulated or downregulated in protoplasts 4h and 24h after transfection as well as protonema (PN) compared to freshly isolated protoplasts (0h). (a) The number of DEGs identified in one haploid line (Haploid A, brown) and in one diploid line (Diploid A, blue). DEGs are combined from identification in the microarray data (filtered for a |log2 fold change| > 1 and p < 0.001) and the SuperSAGE data (filtered for a GFOLD(0.01) value of < −1 or > 1). (b) Overlap between the upregulated genes at 24h in the haploid line with the upregulated genes at 24h in the diploid line. (c) Overlap between the downregulated genes at 24h in the haploid line with the downregulated genes at 24h genes in the diploid line.

Time-dependent DEGs were computed with samples taken from protoplasts at 4h or 24h after transfection versus protoplast samples taken directly after isolation (0h), separately for the haploid and diploid lines in microarray and SuperSAGE data, respectively (Supplementary Table T6). In both the haploid and diploid lines there was no extensive overlap between the DEGs from the second microarray and SuperSAGE experiment showing the advantage of combining both methods (Supplementary Figure S3). We observed for the haploid WT 1148 DEGs at 4h and 4000 DEGs at 24h (SuperSAGE data) and for the line Haploid A 3142 DEGs at 4h and 4663 DEGs at 24h (combination of DEGs from microarray and SuperSAGE data). This makes a combined set of DEGs from both haploid lines with 3823 DEGs at 4h and 6698 DEGs at 24h (467 DEGs at 4h and 1965 DEGs at 24h, respectively, were found in both haploid lines, Supplementary Figure S3). Considering the total number of Physcomitrella genes (32,458 in Physcomitrella V3.3, Fernandez-Pozo et al., 2020; Lang et al., 2018) 3.54% and 12.32% of them were differentially expressed in the WT at 4h and 24h, respectively, while it was 9.68% at 4h and 14.37% at 24h in Haploid A. For Diploid A, we received from microarray and SuperSAGE a set of in total 1628 DEGs at 4h and 4922 DEGs at 24h, being 5.02% and 15.16%, respectively, of the Physcomitrella genes. According to the GO analysis of biological process terms in our initial microarray time series of WT protoplasts (Fig. 1c) we identified in the subsequent microarray data of Haploid A an enrichment of genes encoding enzymes involved in photosynthesis at 24h and additionally at 4h, but they were not enriched in the data of Diploid A. However, in the SuperSAGE data of both haploid and diploid lines genes encoding enzymes involved in photosynthesis were enriched at 24h as well as at 4h (Supplementary Excel sheet 2). In the comparison of 24h versus 0h protoplasts, the percentage of downregulated DEGs was noticeably lower in microarray and SuperSAGE measurements of the diploid line than in the haploid lines (Fig. 2, Supplementary Table T6). For the haploid lines the percentage of downregulated DEGs at 24h was in Haploid A 39.95% and 73.66% in microarray and SuperSAGE data, respectively, and 70.83% in the SuperSAGE data of WT whereas for Diploid A 16.84% of the DEGs at 24h were downregulated in the microarray data and 26.44% in the SuperSAGE data. In each of the analysed data sets more than half of the upregulated DEGs at 24h in the haploid lines were also upregulated in the diploid line (e.g. 72.17% for Haploid A; Fig. 2b). Similarly, most of the downregulated DEGs at 24h in the diploid line were also downregulated in the haploid lines (e.g. 80.05% compared to Haploid A; Fig. 2c).

### Combination of microarray and SuperSAGE sequencing yields a high amount of new DEGs

Xiao et al. (2012) investigated the transcriptomic responses in regenerating Physcomitrella protoplasts and their reprogramming into stem cells, comparing gene expression with digital gene expression tag profiling (DGEP) at four different time-points: 0h, 24h, 48h and 72h. We looked for similarities and differences in the DEGs identified from the 24h versus 0h comparison in that study and the DEGs computed from protoplast samples from 24h after transfection versus 0h after isolation using the combined set of microarray and SuperSAGE data from two haploid lines (WT, Haploid A) in our current study (Supplementary Table T6). Of the 1195 DEGs at 24h versus 0h from Xiao et al. (2012) that were translated into Physcomitrella V3.3 IDs (5 of the 1095 DEGs did not map to any V3.3 ID; 115 mapped to multiple IDs), 801 (69.71%) were also identified in at least one of our datasets of the haploid lines (Supplementary Figure S3), whereas we identified 5,898 additional DEGs. Among the 394 genes that were only found to be differentially expressed by Xiao et al. (2012), some with high log2 fold changes in expression of > 5 or < −5 are for example a homologue to an *Arabidopsis* threonine aldolase (Pp3c4_31180) and cytochrome P450 (Pp3c11_6580). Genes with high expressional changes at 24h that were only found in our data are amongst others a gene encoding a desiccation-related protein of the LEA family (Pp3c17_8560), and an ap2 erf domain-containing transcription factor (Pp3c9_4590). In Xiao et al. (2012), 65.02% of the DEGs were downregulated at 24h versus 0h. According to our SuperSAGE data of both haploid lines at 24h versus 0h these are 70.83% and 73.66%, respectively, while according to our microarray data of Haploid A only 39.95% DEGs were downregulated (Supplementary Table T6).

### Key players of HR and NHEJ are differentially expressed in haploid and diploid lines during protoplast regeneration

In total we identified, manually or via the corresponding biological process GO term, 255 genes that are relevant for DNA repair among the combined set of DEGs from microarray and SuperSAGE data at 4h or 24h after transfection during protoplast regeneration compared to freshly isolated (0h) protoplasts (Supplementary Excel sheet 1). Most of the DNA repair-relevant DEGs that act in HR or NHEJ, the two main DNA-DSB repair pathways, were upregulated. In WT and Haploid A we identified 29 HR-relevant DEGs and a comparable number of 30 HR-relevant DEGs was also detected in Diploid A. 18 of these DEGs occurred in haploid and diploid lines (Supplementary Excel sheet 1). For some differential expression occurred in haploids and diploids at different time-points during protoplast regeneration. For example, the gene encoding the HR-relevant protein Rad50 (Pp3c10_3760; Kamisugi et al., 2012) was upregulated specifically: Whereas upregulation in haploids occurred exclusively at 4h, it occurred in diploids only at 24h. In contrast, we observed at 24h in lines of both ploidies a time-dependent upregulation of REV1 (Pp3c22_4740), a HR-promoting protein (Sharma et al., 2012). Six of the DEGs encode proteins with a role in NHEJ (Supplementary Excel sheet 1) with XRCC4 (Pp3c1_38430), Ku70 (Pp3c18_7140) and Ku80 (Pp3c22_11100) as the key proteins of NHEJ (Weterings and Chen, 2008). The expression of these three genes was upregulated in haploid as well as in diploid lines. The upregulation of XRCC4 was consistent over time at 4h and 24h, whereas the upregulation of Ku70 and Ku80 started only later at 24h. The other NHEJ-related genes were all exclusively upregulated at 24h. These are PRKDC (Pp3c9_15240), POLL (Pp3c15_19010) and ATM (Pp3c2_23700). PRKDC encodes a protein kinase that is recruited to the ends of DNA by the Ku70/KU80 complex (Davis and Chen, 2013). POLL encodes Polymerase λ that plays a role in gap-filling (Lee et al., 2004). ATM encodes a protein kinase that is activated by DNA-DSBs, triggering amongst others the cell-cycle checkpoint signalling and DNA repair (Maréchal et al., 2013). ATM seems to play a role in NHEJ and in HR (Bakr et al., 2015; Weterings and Chen, 2008; Zha et al., 2011) and was the only NHEJ-related DEG that was differentially expressed in the diploid line but not in the haploid lines. Further, we found among the DEGs genes encoding proteins of other DNA-repair pathways like nucleotide excision repair and base excision repair (Supplementary Excel sheet 1).

### Gene expression in diploid and haploid protoplast differs at various time-points

Next, we determined DEGs between haploid and diploid lines separately for freshly isolated protoplasts (0h) and 4h and 24h after transfection in regenerating protoplasts as well as in protonema samples. In the microarray data only few genes with altered expression between haploid and diploid lines were identified (Table 2, column DEGs in microarray; Supplementary Table T7) and the differences in gene expression levels were only small (Supplementary Excel sheet 1).

**Table 2.**
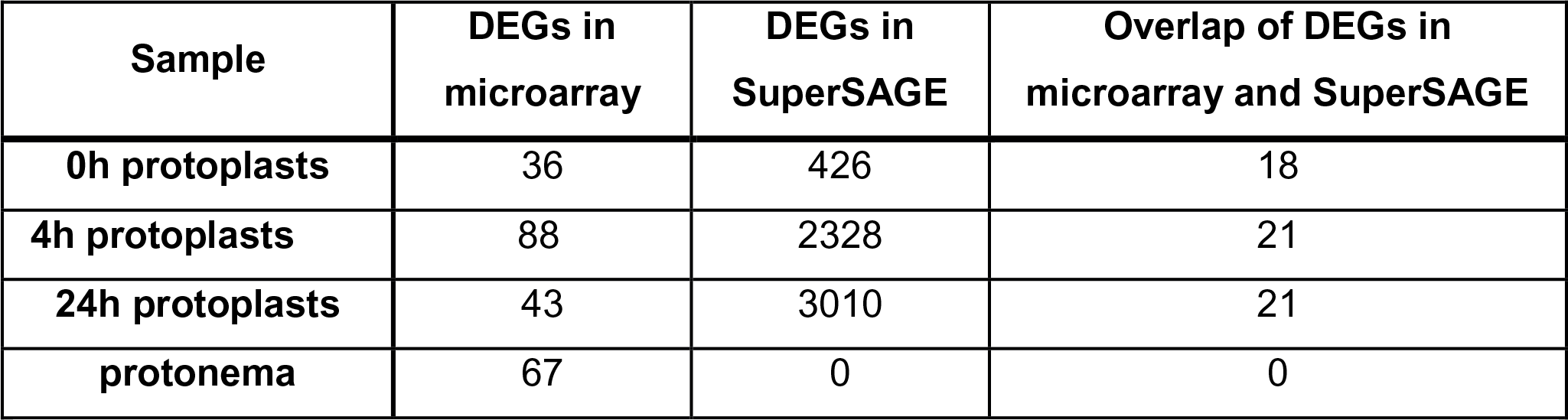
Number of DEGs between diploid and haploid cells at different time-points after transfection. Microarray and SuperSAGE analysis were performed on cells from Diploid A compared to cells from Haploid A. DEGs from the microarray experiment were determined with the Expressionist Analyst Pro software and were filtered for |log2 fold change| > 1 and p < 0.001. The SuperSAGE data analysis was performed with GFOLD and DEGs were filtered for a GFOLD(0.01) value of < −1 or > 1.

| Sample | DEGs in microarray | DEGs in SuperSAGE | Overlap of DEGs in microarray and SuperSAGE |
| --- | --- | --- | --- |
| 0h protoplasts | 36 | 426 | 18 |
| 4h protoplasts | 88 | 2328 | 21 |
| 24h protoplasts | 43 | 3010 | 21 |
| protonema | 67 | 0 | 0 |

Further, we performed pairwise comparisons between haploid and diploid SuperSAGE libraries (Table 1). The pairwise SuperSAGE data analysis of protonema samples from Diploid A versus Haploid A yielded no DEGs while it was 67 in the microarray analysis of the same lines. In contrast, we found more pronounced ploidy-dependent transcriptomic differences in the SuperSAGE data of protoplasts than in the microarray data (Table 2, Supplementary Table T7). There was a moderate overlap between DEGs in microarray and SuperSAGE data (Table 2, Supplementary Figure S4). Most of the DEGs were specific to a certain time-point, while only few of them were common among all protoplast stages (48 in SuperSAGE, 0 in microarray; Fig. 3). Examples are some membrane and surface proteins (e.g., Pp3c3_22320 and Pp3c6_11380) as well as an oxidoreductase (Pp3c4_15600). The same trend that the majority of the DEGs only appeared at one specific time-point occurred also in pairwise comparisons using additionally Diploid B and WT (Supplementary Figure S4, Supplementary Excel sheet 1). There were noticeable differences in the number and identity of DEGs between analyses with different combinations of plant lines (Supplementary Figure S4). An additional principal component analysis (PCA) of the 17 SuperSAGE libraries revealed that the plant material used (protonema or protoplast) and the protoplast regeneration states (0h, 4h, 24h) had the strongest contribution to the variance in gene counts between the libraries, whereas the contribution of ploidy level was inferior (Supplementary Figure S5). Besides, at 4h and 24h there was a visible variance in the gene counts of the two haploid lines.

**Fig. 3.**
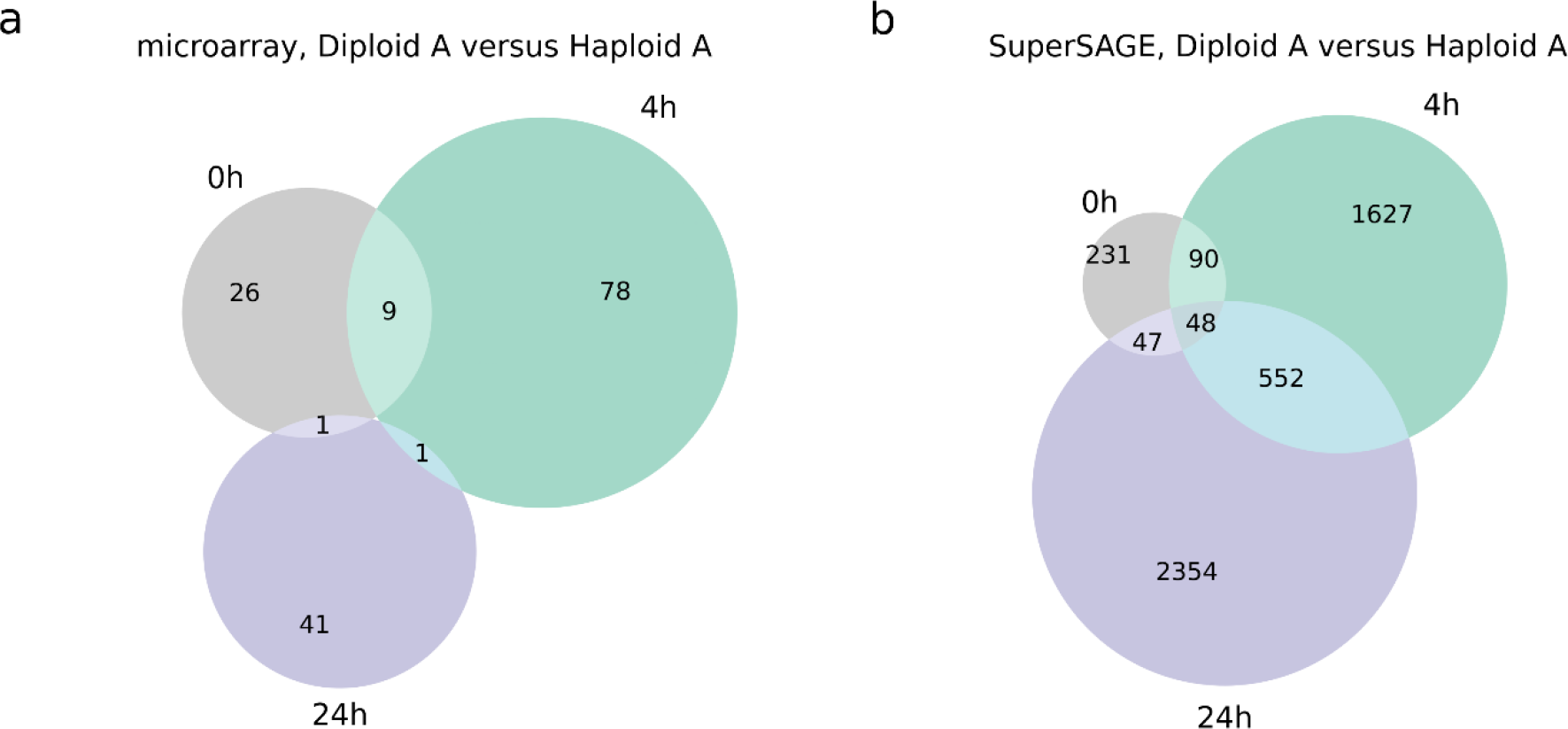
Overlap of DEGs identified from pairwise comparison between one haploid (Haploid A) and one diploid (Diploid A) line at different protoplast stages. DEGs were determined by microarray analysis (a) and SuperSAGE libraries (b) from protoplast samples (grey: freshly isolated protoplasts (0h), green: protoplasts 4h after transfection, purple: protoplasts 24h after transfection. DEGs from the microarray experiment were determined with the Expressionist Analyst Pro software and were filtered for |log2 fold change| > 1 and p < 0.001. The SuperSAGE data analysis was performed with GFOLD and DEGs were filtered for a GFOLD(0.01) value of < −1 or > 1.

### Ploidy affects expression of 3.7% of the Physcomitrella genes

To gain more insight into ploidy-specific gene expression, we performed two analyses for differential expression with a two-factor design. The two-factor design was set up to test for the changes in gene expression caused by ploidy taking the impact of tissue and time into consideration. This design allowed the inclusion of all SuperSAGE datasets in a ploidy-dependent differential gene expression analysis, resulting in a more robust test. The applied tool for these investigations was DeSeq2, which is suitable for multiple-factor analysis with few replicates (Love et al., 2014). For the first analysis, all 17 SuperSAGE libraries from the two haploid lines WT and Haploid A as well as from Diploid A were used, including protoplast libraries from different time-points (freshly isolated protoplasts, 4h and 24h after transfection) as well as protonema libraries (all the libraries are listed in Supplementary Table T3). In a second analysis, we used the same two-factor design, but exclusively compared datasets derived from protoplasts of Diploid A with protoplast of the haploid WT control, omitting the protonema samples to ensure a more homogenous data collection. In the first analysis, we identified 159 ploidy-dependent DEGs (89 upregulated and 70 downregulated in diploid samples). The second analysis yielded 1170 DEGs (635 upregulated and 535 downregulated in diploid samples). Combining the genes with ploidy-dependent expression from both analyses resulted in a total set of 1202 DEGs (127 DEGs were identified in both analyses), comprising 3.7% of the *P. patens* genes according to the latest genome assembly (32,458 in V3.3; Fernandez-Pozo et al., 2020; Lang et al., 2018). The rate of alteration in gene expression was mostly moderate. Particularly high differences in transcript abundances of more than 8 fold (|log2 fold change| >3) were detected only rarely (for 5 upregulated and 1 downregulated genes) and exclusively in the second two-factor analysis (Supplementary Table T8). We found the highest upregulation for a gene encoding a glycosyl-hydrolase-family 88 protein (Pp3c23_940), which cleaves saccharide bonds (Davies and Henrissat, 1995). The gene with the highest downregulation encodes a small subunit ribosomal protein S27a (Pp3c4_19000) that can play a role in disease resistance and cell death (Xu et al., 2019). Other genes with changes of more than 8 fold encode a chalcone-flavonone isomerase 3 related protein (Pp3c4_25770) and a 3-hydroxyisobutyryl-Co A hydrolase (Pp3c22_10130) as well as two unannotated genes (Pp3c4_17790 and Pp3c2_21940). Two of the most prominently downregulated genes from the first two-factor analysis (Supplementary Table T9) encode proteins with reported functions related to cell wall organization: A BR-signalling kinase (Pp3c20_8180; Rao et al., 2017) and an expansin (Pp3c3_16280; Marowa et al., 2016; Schipper et al., 2002).

### Ploidy-dependent expression of genes involved in plant morphogenesis, cell cycle regulation, DNA-DSB repair and DNA accessibility

The observation of an altered phenotype after a WGD reported for several species (Andalis et al., 2004; Cohen et al., 2013) motivated us to look for differential regulation of genes that might contribute to phenotypic alterations between haploid and diploid Physcomitrella lines. In the DEGs from the two-factor analyses we searched for moderately to strongly differentially expressed genes with a |log2 fold change| > 1.5 with a biological process GO term assigned considering development process (GO:0032502), growth (GO:0040007) or any of their child terms. The search yielded in total 9 DEGs (Supplementary Table T10). These include a gene associated with anatomical-structure development (Pp3c1_10860, log2 fold change of 1.85), leaf morphogenesis (Pp3c22_2330, log2 fold change of 1.57) and regulation of meristem growth as well as root hair elongation (Pp3c13_1640, log2 fold change of 1.13 in the first two-factor analysis and 1.56 in the second two-factor analysis).

Subsequently, we performed a targeted search in the pairwise time-point-specific comparisons between the ploidies and the results from the two-factor analyses for differential expression of genes with a role in DNA-DSB repair, cell cycle regulation and DNA accessibility, that might contribute to the higher rates of GT via HR in members of the haploid-dominant bryophytes compared to angiosperms. Considering genes with a direct role in DNA-DSB repair, target genes of interest could be key players for the HR mechanism that have a reduced expression in diploids compared to haploids or in contrast act in an end-joining pathway with elevated expression in diploids. In the combined set of DEGs from the microarray and SuperSAGE data at 4h or 24h after transfection, we observed that most genes belonging to the HR and the NHEJ pathway, respectively, were upregulated in diploid cells compared to haploid cells at one or several time-points after protoplast transfection (Supplementary Excel sheet 1). From the 21 identified HR-relevant DEGs only 3 were downregulated in diploids. These genes encode the ATPase YcaJ (Pp3c23_21270; −1.38 log2 fold change at 4h), DNA polymerase I (Pp3c14_14550; −1.59 log2 fold change at 4h) and poly(ADP-ribose) polymerase (Pp3c8_13220; −1.24 log2 fold change at 4h). Only 3 NHEJ-related ploidy dependent DEGs were identified; all of them being upregulated. They encode ATM (Pp3c2_23700; 1.06 log2 fold change at 24h), the DNA-dependent protein kinase catalytic subunit (Pp3c9_15240; 1.87 log3 fold change at 24h) and XRCC4 (Pp3c1_38430, 1.46 and 1.22 log2 fold change at 4h and 24h respectively). In this context the latter two are known players of NHEJ (Brouwer et al., 2016; Chang et al., 2017; Graham et al., 2016). Additionally, the gene encoding Polymerase Q (POLQ) was upregulated (Pp3c5_12930; 1.46 log2 fold change at 24h). It acts in alternative end-joining and is a potential inhibitor of HR in Physcomitrella (Mara et al., 2019).

We selected candidate genes for validation with qRT-PCR from the list of DEGs in our two-factor analyses. From the previously mentioned genes, XRCC4 was the only one that was also a DEG in one of the two-factor analyses, representing the gene with the strongest transcriptomic difference between haploids and diploids that is directly involved in the repair of DNA-DSBs. In the second two-factor analysis, comparing datasets derived from protoplasts of Diploid A with protoplast of the haploid WT at 4h and 24h, it was upregulated with a log2 fold change of 1.76 in the diploid line. Hence, we chose XRCC4 for validation with qRT-PCR. Furthermore, three additional genes with a function in cell-cycle regulation or DNA accessibility were selected from the upregulated DEGs of the two-factor analyses for validation with qRT-PCR (Table 3): CENPE (Pp3c22_20430), cyclin D2 (Pp3c9_8300) and H3K4-Methyltransferase (Pp3c4_16880).

**Table 3.**
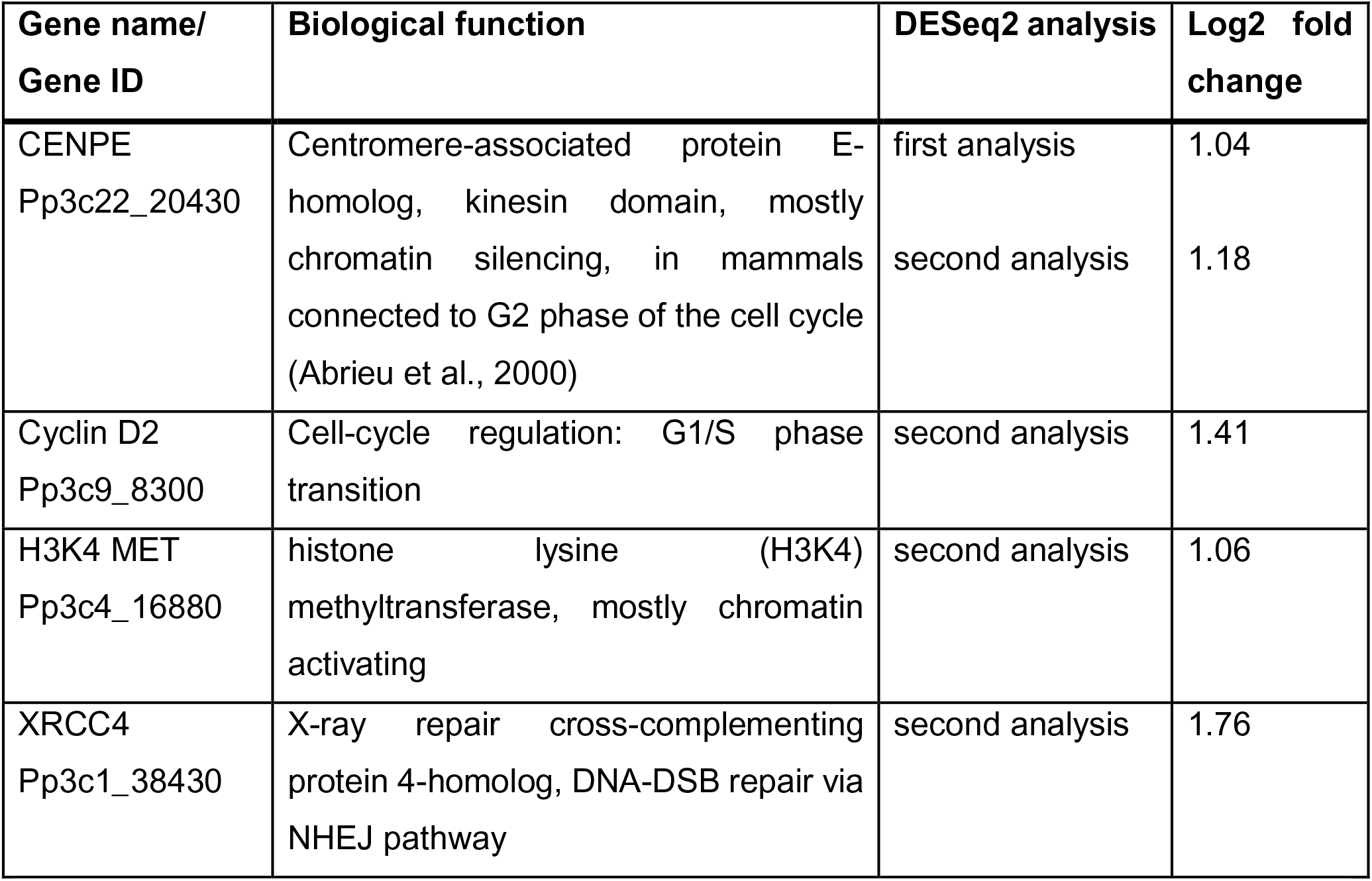
Overview of ploidy-dependent expressed genes with reported functions in DNA-DSB repair, cell cycle regulation and DNA accessibility identified by two-factor analyses. Expression fold changes is given for diploid cells in comparison to haploid cells.

| Gene name/<br>Gene ID | Biological function | DESeq2 analysis | Log2 fold<br>change |
| --- | --- | --- | --- |
| CENPE<br>Pp3c22_20430 | Centromere-associated protein E-homolog, kinesin domain, mostly chromatin silencing, in mammals connected to G2 phase of the cell cycle (Abrieu et al., 2000) | first analysis<br>second analysis | 1.04<br>1.18 |
| Cyclin D2<br>Pp3c9_8300 | Cell-cycle regulation: G1/S phase transition | second analysis | 1.41 |
| H3K4 MET<br>Pp3c4_16880 | histone lysine (H3K4) methyltransferase, mostly chromatin activating | second analysis | 1.06 |
| XRCC4<br>Pp3c1_38430 | X-ray repair cross-complementing protein 4-homolog, DNA-DSB repair via NHEJ pathway | second analysis | 1.76 |

### qRT-PCR validates upregulation of XRCC4 in diploid protonema cells

To experimentally validate these DEGs, transcript abundances in three lines (WT, Haploid A, Diploid A) were quantified via real-time qRT-PCR. RNA was isolated from protonema in biological and technical triplicates. In the qRT-PCR analysis, an upregulation of H3K4-Methyltransferase and cyclin D2 as observed in the first two-factor analysis (Diploid A versus WT and Haploid A including all protoplast and protonema samples) as well as of CENPE as observed in the first and second two-factor analyses (Diploid A versus WT including all protoplast samples) was not supported in diploid protonema (Fig. 4). In contrast, a ploidy-dependent expression of XRCC4 in protonemal tissue was validated as noticeably different from WT control in the diploid line by the qRT-PCR approach with a 1.86±0.16 log2 fold increase in transcript abundances for Diploid A. Next, we checked XRCC4 expression in natural developmental stages of WT Physcomitrella in publicly available datasets as compiled by Ortiz-Ramírez et al. (2016). We selected the four datasets that contain values for the gene expression in the sporophyte from Gransden or Reute ecotype: RNAseq developmental stages v3.3 (including data from Reute and Gransden ecotype), CombiMatrix Developmental stages gmv1.2 (including data from Reute and Gransden ecotype), NimbleGen Developmental and Mycorrhiza gmv1.6 (including data from Reute ecotype; mycorrhiza exudate and heat treated samples were not considered) and NimbleGen gmv1.6 (including data from Gransden ecotype). The first three of these datasets contained sporophytic data only from the Reute ecotype, whereas in the last it was from the Gransden ecotype. In all the datasets, we observed a tendency for high XRCC4 expression in the natural diploid sporophytic developmental stages, compared to the haploid stages (Supplementary Figure S6). The highest XRCC4 expression level was always in one of the sporophytic stages even though XRCC4 expression level was in some sporophytic developmental stages of the datasets lower or comparable to other haploid stages. This was the case in the NimbleGen dataset, where only the brown sporophyte of the Gransden ecotype showed especially high XRCC4 expression levels while in the earlier green sporophytic stages XRCC4 expression was not considerably enhanced compared to archegonia and spores (Supplementary Figure S6c). In contrast, in the NimbleGen Developmental and Mycorrhiza dataset and the RNAseq developmental stages dataset XRCC4 expression was highest in the green sporophyte of the Reute ecotype but the level in brown sporophytes was lower and comparable to that in juvenile gametophores (Supplementary Figure S6b, d). Similarly to XRCC4, the highest expression level of cyclin D2 in all four datasets was in one of the sporophytic stages (Supplementary Figure S7a-d). However, cyclin D2 was also strongly expressed in several other developmental stages, for example in the spores (ecotype Gransden) in the RNAseq developmental stages dataset (Supplementary Figure S7d). For CENPE and the H3K4-Methyltransferase, the expression in the sporophytes was not considerably higher or even much lower than in the other developmental stages (Supplemental Figs. S8, S9). Only the embryo data of the NimbleGen dataset from Ortiz-Ramírez et al. (2016) showed a quite high expression of both genes, but the expression drastically decreased in the later developmental stages of the sporophyte (Supplemental Figs. S8c, S9c).

**Fig. 4.**
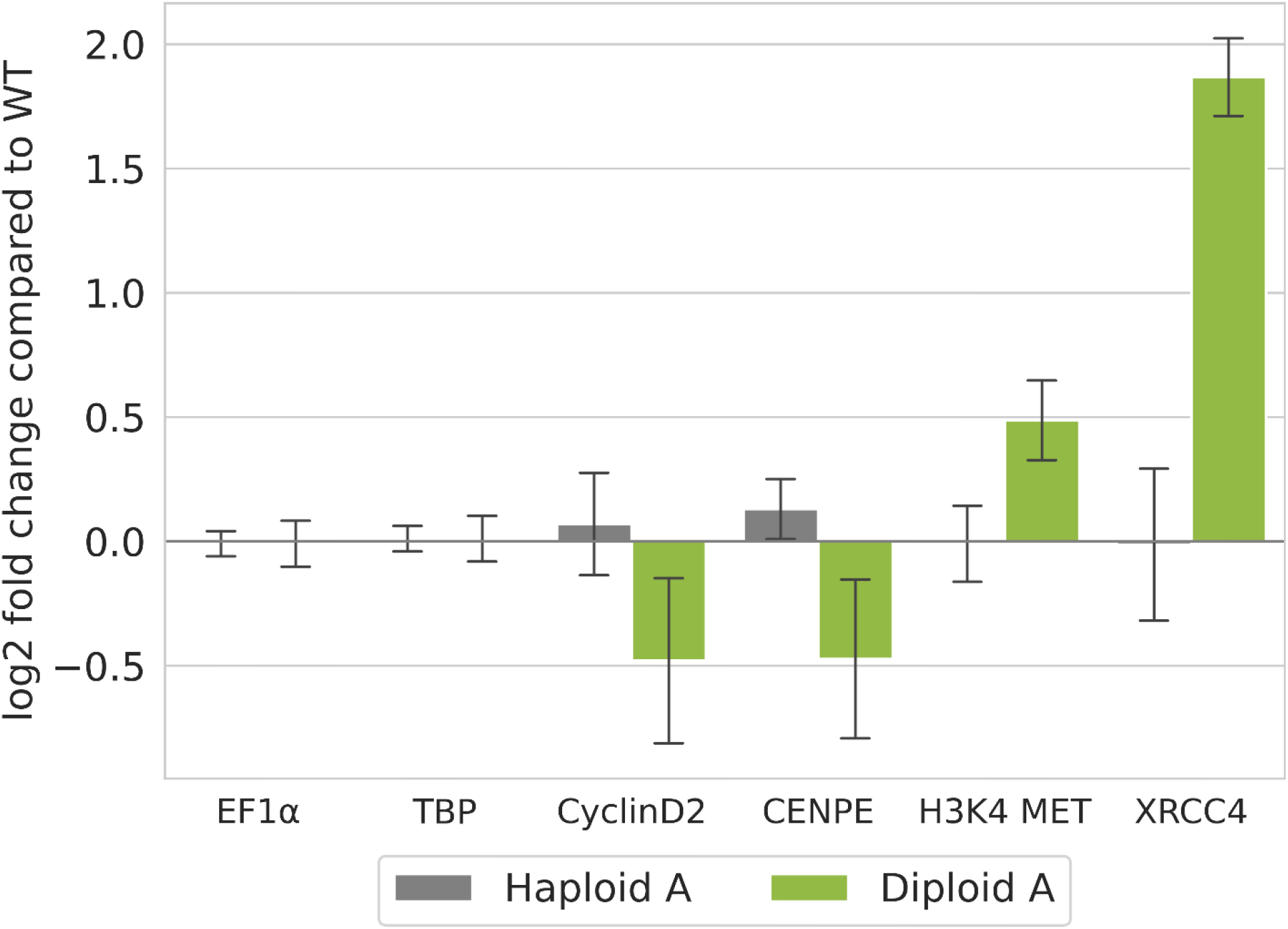
XRCC4 is upregulated in protonemata of the diploid Physcomitrella line. Relative transcript abundance in haploid and diploid Physcomitrella lines compared to WT as measured by real-time qRT-PCR. Normalized relative quantities were computed for each of three biological replicates according to Hellemans et al. (2007). Depicted is the mean log2 fold change over the replicates, error bars represent the standard deviations. EF1α and TBP are shown as reference for ploidy-independent gene expression.

### GT rate is reduced in diploid Physcomitrella cells

In order to investigate if the DNA-DSB repair is influenced by ploidy level, the rates of GT via the HR pathway were quantified in two haploid lines (Haploid A, Haploid B) and three diploid lines (Diploid A, Diploid B, Diploid C). For this purpose, the lines were transformed with a KO construct containing a 1920 bp cDNA fragment encoded by the cold-responsive gene Pp3c21_180V3, encoding sphingolipid fatty acid desaturase (PpSFD) (Beike et al., 2015; Hohe et al. 2004; Resemann et al., 2021). The GT rate was defined as the number of KO plants divided by the total number of transformants – the latter characterized by survival on selection medium and the presence of the selection marker (npt II; verified via PCR analysis). All transformants were tested by flow cytometry to determine their ploidy. This procedure resulted in the identification of 244 haploid and 302 diploid transformants (Fig. 5b). All 546 plants were tested with primers JMLKO25L and JMLKO25R flanking the insertion site of the KO cassette (Supplementary Table T1; Supplementary Figure S2). This approach screens for the presence or absence of the intact WT locus of the target gene. KO plants were characterised by the interruption of the WT locus for haploid plants and by interruption at both chromosomes for diploid plants (Fig. 5a). This procedure showed a disruption of the WT locus in 96 haploid KO plants, corresponding to a GT rate of 0.39. In diploids, successful GT requires the knockout in both chromosomes and hence the expected GT rate would be the square of the GT rate in haploid plants: 0.39 ∗ 0.39 = 0.15. However, for the diploid lines only 16 plants had a disrupted WT locus on both chromosomes, corresponding to a GT rate of 0.05. This value is significantly lower than the predicted rate of 0.15 (Fisher’s exact test, p < 0.001), revealing that the frequency of GT is significantly reduced in diploid plants. Transformants with successful GT were checked for correct 5’ and 3’ integration of the construct into the Physcomitrella genome via PCR. 51 of the 96 haploid plants showed proper integration patterns on both ends (5’ and 3’), while it was 4 out of 16 in diploids.

**Fig. 5.**
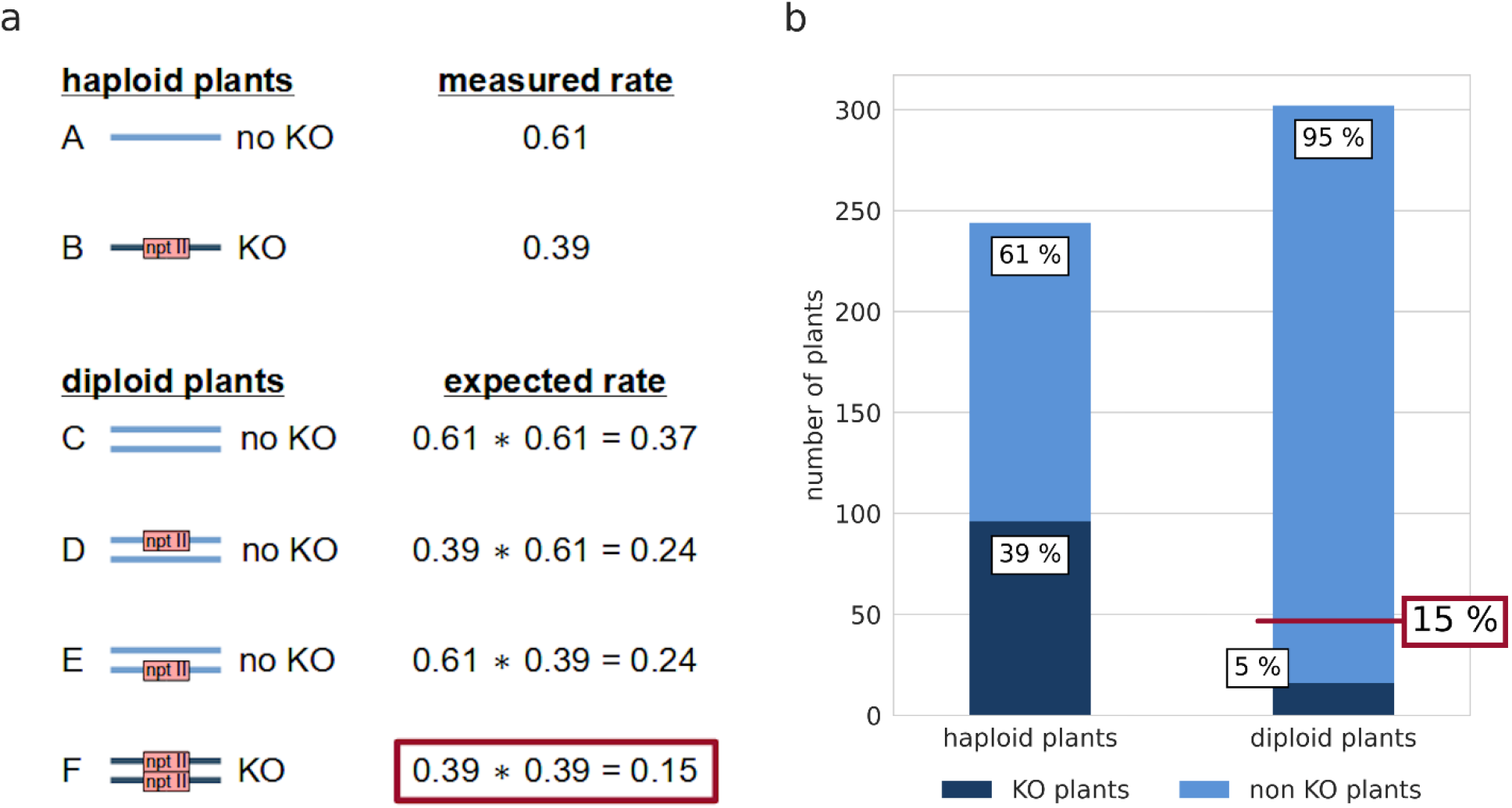
Diploid Physcomitrella cells show significantly lower GT rates. a: Estimation of the expected GT rate in diploid Physcomitrella plants computed from the measured GT rate in haploid plants. Shown are the observed rates of untransformed (A) and transformed (B) haploid plants as well as the expected rates of diploid plants having no integration of the cDNA construct in both chromosomes (C), having the cDNA construct integrated in only one chromosome (D and F) and having a full knock-out (KO) of the target locus in both chromosomes (F). The genomic loci are represented as solid lines and integration of a cDNA-construct containing the npt II cassette as selection marker is indicated. b: Comparison of GT rates in haploid and diploid Physcomitrella plants as determined via PCR analysis. For haploid plants, 96 out of 244 transformants are targeted KOs while for diploid plants, only 16 out of 302 transformants are targeted KOs on both chromosomes. The expected value of targeted KOs under the assumption of equal GT frequency for haploid and diploid plants is 15 % (marked in red).

## Discussion

In this study, we aimed to investigate if a WGD, induced by protoplast fusion, leads to qualitative alterations in gene expression. Such a qualitative change in gene expression might contribute not only to phenotypical changes (Egener et al., 2002; Schween et al., 2005b) but it might also be relevant for diverging GT success rates via HR between angiosperms and members of haploid-dominant bryophytes like the moss Physcomitrella. Here, we used Physcomitrella as a model organism to study differences in the gene expression pattern in haploid and artificially generated diploid plants. Physcomitrella is a particularly suitable model organism for this purpose because it grows in a life cycle where the haploid gametophytic stage is dominant and consequently diploidized Physcomitrella plants have, contrary to wild-type diploid species, two identical copies of each chromosome (autopolyploids). Thereby, we can exclude that observed transcriptomic differences arise from natural variability in the pair of homologous chromosomes, as is the case in allopolyploid hybrids. Here, we designed a multi-layer study using at first a fine-grained microarray time-series from samples taken during the regeneration period of transfected haploid WT protoplasts, ranging from the time-point of protoplast isolation to 72h after addition of foreign DNA, to investigate how protoplast recovery affects the gene expression in the time period when GT is expected to take place. This first time-series was followed by a second microarray experiment and the more sensitive SuperSAGE technology to identify general transcriptomic differences between haploid and diploid lines at three selected time-points during protoplast regeneration and to subsequently concentrate on relevant genes. Finally, we set up two-factor analyses to test for the changes in gene expression caused by ploidy taking into consideration the impact of protoplast and protonema tissue as well as the time-point.

For protoplasts, the removal of the cell wall and the isolation of the single cell exposing it to a new environment represent massive, highly stressful interventions. During regeneration, Physcomitrella protoplasts partially reprogram into stem cells. Xiao et al. (2012) identified transcriptomic changes during reprogramming via digital gene expression tag profiling (DGEP), employing three comparisons between different time-points after protoplast isolation: 24h versus 0h, 48h versus 24h and 72h versus 48h. Here, we investigated the process via a microarray time series and concentrated on early events. Therefore, we sampled tissues during the first 24 hours after transfection, using protoplast samples from 1h, 4h, 6h, 24h, and additionally from 72h after transfection, and compared the gene expression to freshly isolated (0h) protoplasts.

At 1h, protoplasts had apparently not yet adapted gene transcription levels to the changed situation, as we found no DEGs compared to freshly isolated protoplasts. In contrast, at 4h cells had already started reprogramming, as evidenced by more than 200 DEGs. These encode, among others, enzymes involved in L-phenylalanine catabolism or in aromatic amino acid family catabolism. In our study, the phenylalanine ammonia lyase, an enzyme that catalyzes the first step of the phenylalanine catabolic process (Hyun et al., 2011), was differentially expressed at 4h and was also among the DEGs at 72h. This gene is regulated in reaction to biotic and abiotic stresses and is the entry-point enzyme of the phenylpropanoid pathway, thereby supplying the basis for the synthesis of many downstream products like flavonoids. These compounds can function as UV-filter, antioxidant agents and in drought resistance (Samanta et al., 2011). Moreover, plant flavones are important extracellular signals for the root microbiome, especially under nitrogen deprivation (Yu et al., 2021). Further at 4h, among others DEGs with the GO term glycolytic process were enriched (Fig. 1c). Among the DEGs identified in our study at 4h was the glycolysis key player glyceraldehyde-3-phosphate dehydrogenase (Supplementary Excel sheet 1). This enzyme is inhibited under oxidative stress by a redox modification of a cysteine residue. Additionally, there is evidence that in plants the glyceraldehyde-3-phosphate dehydrogenase fulfils non-metabolic functions under stress conditions supported by redox modifications of the enzyme (Gurrieri et al., 2021; Schneider et al., 2018, Wood et al., 2004). Nearly the whole set of genes that were differentially expressed at 6h also belongs to the group of genes that plays a role in the process of protoplast regeneration at 4h as shown by a high overlap of DEGs between 4h and 6h (Fig. 1b).

A second peak became apparent at 24h, resulting in the highest observed number of DEGs. At this time-point, an enrichment of DEGs with a function in aminomethyltransferase activity were apparent in our data, which is in accordance with Xiao et al. (2012). In contrast to those authors, we did not detect enriched GO terms considering protein folding or reaction processes to the environment like response to salt stress, cold or heat at 24h. In accordance to Xiao et al. (2012), however, we observed a significant enrichment of DEGs acting in photosynthesis at 24h. In the study of Xiao et al. (2012) most of the photosynthesis-related DEGs were downregulated at this time-point. A reduction of photosynthesis is a response to both abiotic and biotic stress and is assumed to represent a trade-off in the distribution of means between growth and defense (Attaran et al., 2014; Cohen and Leach, 2019). In Xiao et al. (2012) the expression of many photosynthesis-related genes increased again at 48h compared to 24h, probably to supply the necessary energy for the protoplasts that had re-entered the cell cycle at this time-point.

Accordingly, three days after protoplast isolation (72h) the DEGs identified in our study were less dominated by genes associated with photosynthetic processes, indicating that the energy management of the cell started to return to the normal state. At this time, most of the Physcomitrella protoplasts have undergone the first cell division (Xiao et al., 2012), arriving at a new stage with new transcriptomic requirements. Accordingly, we observed more than 200 new DEGs indicating the re-initiation of various physiological and cell cycle dependent processes stopped upon protoplast isolation.

Overall, protoplast regeneration requires the regulation of important cellular processes that change gradually over time, and only 18 genes were always differentially expressed between 4h and 72h. Among them is a gene of the GRAS family encoding a homologue to the gibberellin regulatory protein RGL1 of *Arabidopsis*. In Physcomitrella the gibberellin precursor ent-kaurene plays a role in developmental regulation (Hayashi et al., 2010) while the gibberellin signaling pathway controlling plant development as it exists in angiosperms seems not to be present in Physcomitrella (Vandenbussche et al., 2007). Another gene that was differentially expressed at all time-points encodes a catalase. Catalases degrade H_2_O_2_, and are indicators for oxidative stress (Smirnoff and Arnaud, 2019; Yong et al., 2017). Upregulation of various types of stress-response genes during cell wall regeneration of protoplasts is reported in cotton and rice (Sharma et al., 2011; Yang et al., 2008).

In a next step, we generated more detailed transcriptomic data from several haploid and diploid lines using microarray and additionally SuperSAGE technology. We analysed freshly isolated protoplasts and protoplasts at 4h and 24h after transfection, respectively. The DEGs of the haploid lines identified in our SuperSAGE and microarray data at 24h compared to freshly isolated protoplasts covered 69.71% of the 1195 identified DEGs (translated from gene model version V1.6 to V3.3, see results) at this time-point in regenerating haploid Physcomitrella protoplast from Xiao et al. (2012). In addition, we identified 5898 DEGs, presumably due to different search criteria: We used different detection methods (microarray, SuperSAGE), two different haploid lines, and different filter criteria for DEGs (|log2 fold change| > 1 and p < 0.001 for microarray data and GFOLD(0.01) value < −1 or > 1 for SuperSAGE data in our studies; |log2 fold change| ≥ 2, p ≤ 0.01 and FDR < 0.01 in Xiao et al. (2012)). Examples of DEGs with high fold changes that were not common between our study and that from Xiao et al. (2012) are cytochrome P450 in Xiao et al. (2012) and an ap2 erf domain-containing transcription factor in our data, both playing roles in plant development and stress responses (Gu et al., 2017; Hiss et al., 2014; Xu et al., 2015).

Next, we compared haploid and diploid protoplasts during reprogramming and regeneration considering changes in gene expression at 4h and at 24h after transfection versus protoplasts directly after isolation. A PCA of the SuperSAGE libraries indicated that a WGD had a much weaker effect on gene expression than the generation of protoplasts itself, or their subsequent regeneration. Indeed, the diploid lines seemed to follow some similar time-steps of protoplast regeneration as observed in haploids. In all analysed samples of diploid and haploid lines more genes were differentially expressed at 24h than at 4h (Supplementary Table T6) and independent of ploidy level, genes with a role in photosynthesis were enriched in the DEGs of the SuperSAGE data at 4h and 24h (Supplementary Excel sheet 2). In lines of either ploidy, the important role for DNA-DSB repair in regenerating protoplasts was reflected by differential expression of genes attributed to both types of repair pathways: Genes acting in HR and genes relevant for the NHEJ pathway, most of them being upregulated. For example, in haploids and diploids we detected at different time-points an upregulation of the HR key player Rad50 as well as a synchronous upregulation over time of the NHEJ key player XRCC4.

However, when directly comparing gene expression between haploid and diploid lines using freshly isolated protoplasts, and protoplasts at 4h and 24h after transfection, we discovered several ploidy-dependent DEGs. Surprisingly, most of them were only differentially expressed in diploid versus haploid cells at one of the analysed time-points, hinting towards time-point-specific variations of some processes or pathways in diploids (Table 2, Fig. 3). According to our two-factor analyses on SuperSAGE libraries, in total 3.7% of the Physcomitrella genes were DEGs in response to the WGD, but mostly with only a moderate change in expression. This clearly indicates a ploidy-dependent gene expression pattern in Physcomitrella protoplasts. The reported values of DEGs from diverse plant species after an artificial autopolyploidization varies strongly. For example, the amount of DEGs in diploid versus tetraploid *Paspalum notatum* with 0.49% reported is considerably lower than our observed value of 3.7% in Physcomitrella, whereas in *Zea mays* the reaction to the WGD is stronger with as much as more than 26% DEGs (reviewed in Spoelhof et al., 2017).

The transcriptomic differences between diploid and haploid plants suggest that diploidization might have an influence on the phenotype. Several studies addressed this issue in Physcomitrella. One of them considered in total 500 mock-transformed haploid and diploid plants (Schween et al., 2005b), while two others included data from large mutant collections of 16,203 (Egener et al., 2002) or 73,329 (Schween et al., 2005a) haploid and polyploid transformants. In these studies only less than half of the polyploid plants showed normal growth on (minimal) Knop medium. On the contrary, for the vast majority of the haploid plants studied, as well as in Schulte et al. (2006), who investigated 51,180 haploid knock-out mutants, growth on Knop medium was normal. For example, in Schween et al. (2005b) more than 90% of the haploids grew normally on Knop medium but only 20% of the diploids. Besides, a weak correlation of 0.55 between ploidy and growth on Knop medium was reported. However, considering the growth on Knop medium, the diploid lines analysed in our study were specifically selected not to behave significantly different from the haploid lines. Another feature that correlated with the ploidy level in Schween et al. (2005b) was the rate of coverage with gametophores that was comparable to the wild type in 74.9% of the haploid plants but only in 7.6% of the diploids. Furthermore, the leaf shape correlated with ploidy level and 25% of the diploids had a double leaf tip compared to only 0.1% of the haploids. Multiple phenotypic deviations were a feature correlating with ploidy that happened in about a quarter of the diploids but in none of the haploids. In total, much more haploid than diploid gametophores looked similar to the wild type with 93.1% and 26.3%, respectively. Other features apparently interlinked with the ploidy level as reported in Schween et al. (2005a) are the plant structure and the uniformity of leaves. We identified 9 ploidy-dependent DEGs with a |log2 fold change| in expression of more than 1.5 that are, according to their biological process GO terms, associated with developmental process or growth (Supplementary Table T10). These include genes associated with anatomical-structure development, leaf morphogenesis and regulation of meristem growth.

Next, we searched for ploidy-dependent DEGs associated with DNA-DSB repair, cell cycle regulation and DNA accessibility which is closely associated with the first two processes. Since HR takes place in the same period as protoplast regeneration, survival and regeneration of the protoplast are responsible for a strong “background noise” of DEGs, that might mask subtle transcriptomic responses leading to the homologous integration of a transgene. In our study of diploid versus haploid protoplasts we detected an upregulation of genes required for HR as well as for NHEJ at some time-points. Only few HR-relevant genes were downregulated in diploids, for example DNA polymerase I. Further, the gene encoding Polymerase Q (POLQ) that is unfavourable for GT via HR in Physcomitrella (Mara et al., 2019), was upregulated in the same cells. One especially interesting candidate was the gene encoding the Physcomitrella homologue of XRCC4, a key player in the NHEJ DNA-DSB repair pathway in mammals (Chang et al., 2017), which is also upregulated in Physcomitrella after bleomycin-induced DNA damage in haploid cells (Kamisugi et al., 2016). Here we detected an upregulation of XRCC4 not only in diploid versus haploid protoplasts at 4h as well as 24h after transfection using the microarray time series and SuperSAGE data, but it was also the DNA-DSB repair-relevant gene with the highest change in expression in the two-factor analysis of the SuperSAGE data (Supplementary Excel sheet 1). Quantitative real-time PCR validated that XRCC4 level is ploidy-dependent with a much higher transcript abundance in diploids than in haploids (up to 2.58±1.0 log2 fold increase in diploid protonema, Fig. 4). An investigation of the XRCC4 expression profile during the Physcomitrella life cycle in publicly available transcript data (PEATmoss; Fernandez-Pozo et al., 2020) revealed a tendency to higher XRCC4 transcript levels in the natural diploid life stage, the sporophyte, than in haploid protonema and protoplasts (Supplementary Figure S6a-d). These findings suggest that the choice of the DNA-DSB repair pathway is most likely dependant on the ploidy level, not only in plants before and after a WGD, but also in the natural haploid and diploid developmental stages of WT Physcomitrella. The existence of such an interdependency between ploidy level and the repair pathway choice was reported for haploid and diploid yeast cells under DNA replication stress (Li and Tye, 2011).

After having identified some DEGs relevant for the repair of DNA-DSBs, and thus potentially for the choice between HR and NHEJ, in haploid versus diploid protoplasts, we analysed GT frequencies between both cell types. The GT construct pRKO25.2 (Hohe et al., 2004) contains a 1920 bp cDNA fragment encoded by the cold-responsive gene Pp3c21_180V3, encoding sphingolipid fatty acid desaturase (PpSFD) (Beike et al., 2015; Resemann et al., 2021) and was utilized to transform two haploid and three WGD lines of Physcomitrella. Surprisingly, transformation of diploid protoplasts yielded only 5% true knockouts instead of the theoretically expected 15% (Fig. 5), revealing a significant suppression of GT after WGD. Hohe et al. (2004) compared gene targeting rates of different single cDNA constructs with mixes of 5 or 10 cDNAs and found no difference in gene targeting rates between single and mixed cDNA, indicating that for the transformation protocol used the uptake of cDNA during transformation is not a limiting factor for homologous recombination. Martin et al. (2009) showed that the production of double FtsZ-mutants can be as effective as the production of single mutants, confirming that the amount of cDNA during transformation is sufficient for several loci at the same time. Hence, in diploid Physcomitrella lines increased expression of the gene encoding XRCC4 correlates with a suppression of GT and thereby, the NHEJ pathway gains in significance over HR, the main DNA-DSB repair mechanism of the haploid-dominant moss (Kamisugi et al., 2006). We interpret high NHEJ rates in diploid lines as a reduced selective pressure for accurate DNA repair via the HR pathway due to the additional information back-up available in form of a second set of chromosomes. Elevated NHEJ rates in diploids support the hypothesis that the haploid phase of Physcomitrella is interlinked with high integration rates of transgenes via HR (Schaefer and Zrÿd, 1997). Yet, ploidy is unlikely the sole factor that determines GT rates in plants for several reasons: i) GT frequencies of seed plants did not increase with haploid tissues (Mengiste and Paszkowski, 1999), ii) GT in other haploid species like *Volvox* is not as efficient as in Physcomitrella (Reski, 1998b), and iii) the GT rate we measured in diploid Physcomitrella plants is still a multiple factor higher than GT rates observed in polyploid angiosperms. Another factor potentially contributing to the GT efficiency in Physcomitrella is the G2/M-phase arrest of the protonema tissue used for transformation. This was, however, unchanged after WGD in our diploid lines.

The differences in gene expression between the analysed haploid and diploid lines having an identical, albeit duplicated, genotype might be to some extent caused by ploidy-dependent epigenetic regulation of the transcriptome. Epigenetic regulation of chromatin accessibility is partially mediated via chromatin marks by methyl or acetyl groups. Xiao et al. (2012) showed that various homologues of methyltransferases are DEGs during protoplast regeneration in Physcomitrella, coinciding with the expected time HR takes place. This may indicate an important mechanism for epigenetic regulation of DNA repair pathways. Indeed, epigenetic alterations (Wolffe and Matzke, 1999) as well as the adaption of gene-regulatory networks and direct changes in the genome structure, among others by an altered transposable element activity or homologous and non-homologous recombination (Adams and Wendel, 2005; del Pozo and Ramirez-Parra, 2015; Liu and Wendel, 2003; Otto, 2007), were reported already to happen in the first generations very shortly after a genome-duplication event. They are a reaction to challenges arising in newly formed polyploids, like genetic instability (Soltis et al., 2015), an increased demand of energy and a higher number of chromosomes to deal with during mitosis (del Pozo and Ramirez-Parra, 2015; Doyle et al., 2008).

With the creation of artificial diploid Physcomitrella plants, we have imitated a WGD event, which is an important driving force of evolution that happened several times over the past 200 million years in land plants (Renny-Byfield and Wendel, 2014; Soltis and Soltis, 2016; van de Peer et al., 2017). Our studies provide an insight into the adaption of gene expression in the decades following a WGD. Such findings might help to retrace how autopolyploids established during evolution. Additionally, with our findings we are one step closer to unmasking the mysteries surrounding GT in plants by further elucidating the regulation of DNA repair mechanisms. Understanding the mechanism of HR is the basis for transferring the technique and efficiency to create genetically modified organisms via GT from Physcomitrella to other plant species (Collonnier et al., 2017). Even with the current advances in GT using CRISPR/Cas systems, they still have limitations and success rates in crops are still low (Barone et al., 2020; Manghwar, et al., 2019). Here, our findings might help in establishing a procedure for reliable introduction of specific traits into crop plants via GT that is an important objective of many plant biotechnological studies and of interest to agriculture.

## Author contributions

C.R. designed and performed research, analysed data and wrote the manuscript. G.W. designed and performed research, and analysed data. G.S. designed and performed research and wrote an initial version of the manuscript. K.L.K. and J.M.L. performed research and analysed data. R.H. performed research and acquired funding. E.L.D. designed research and wrote the manuscript. R.R. designed research, wrote the manuscript and acquired funding. All authors discussed data and approved the final version of the manuscript.

## Acknowledgments

We gratefully acknowledge funding by the Deutsche Forschungsgemeinschaft (DFG, German Research Foundation) under Germany’s Excellence Strategy EXC-2189 (CIBSS to R.R.), the Federal Ministry of Education and Research BMBF (GABI-PRECISE 0315057D to R.H. and R.R.) and the Ministry of Science, Research and Art of the Federal State of Baden-Württemberg (MWK) as part of the Science Data Center funding program BioDATEN. We thank Tanja Egener-Kuhn, Annette Hohe and Anja Martin for initial experiments, Agnes Novakovic for excellent technical assistance and Anne Katrin Prowse for language editing.

## Author statement

The authors declare no competing interest.

## Supplement

**Supplementary Figure S1.**
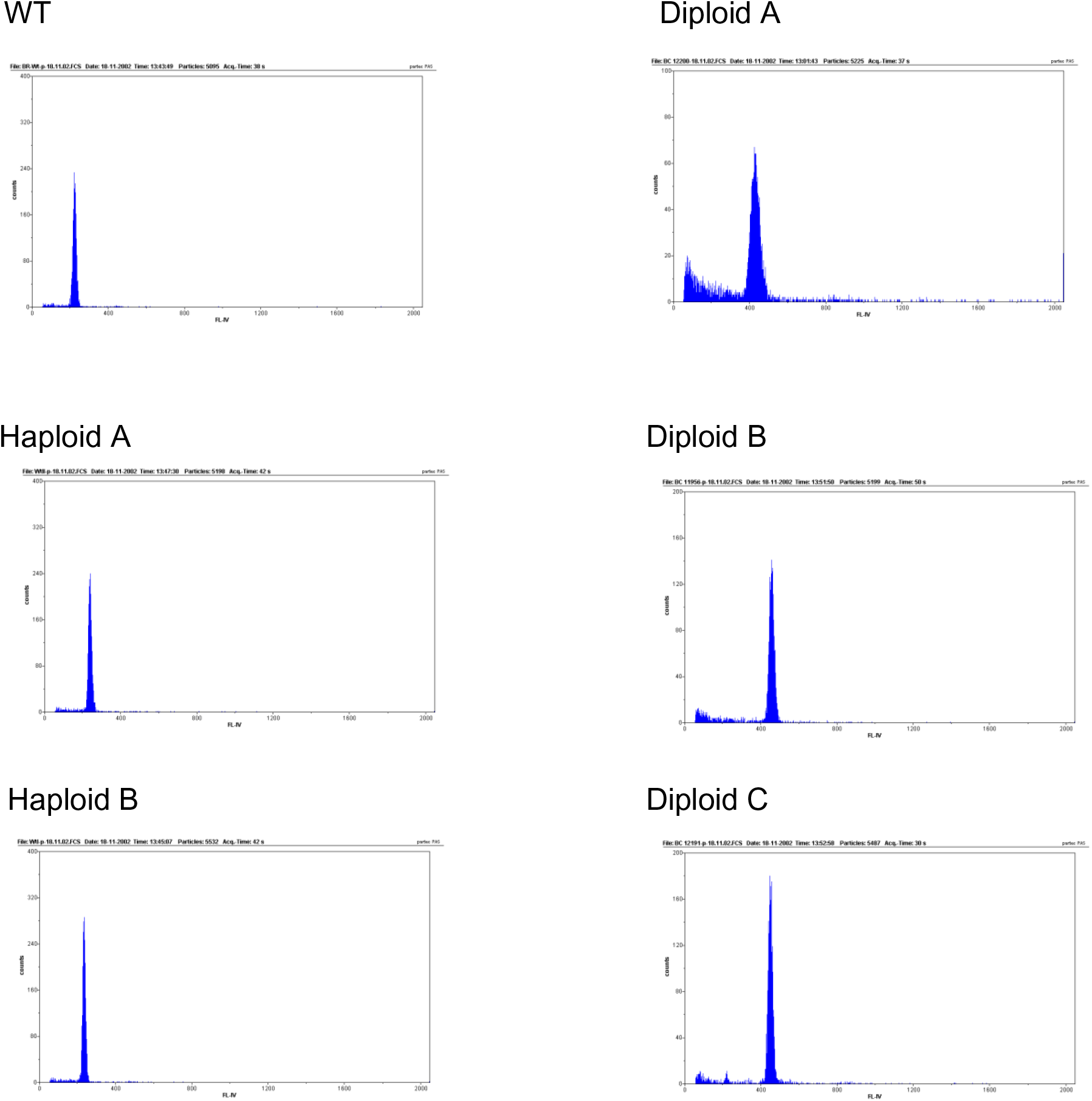
Flow cytometric analyses of haploid and diploid Physcomitrella lines as published in Schween et al. (2005b). For haploid lines (left), peaks at a fluorescence intensity of about 200 represent cells in the G2 phase whereas cells in the G1 phase have a fluorescence intensity of about 100. For diploid lines (right), the peaks at a fluorescence intensity of about 400 represent cells in the G2 phase and cells in the G1 phase have a fluorescence intensity of about 200.

**Supplementary Table T1.**
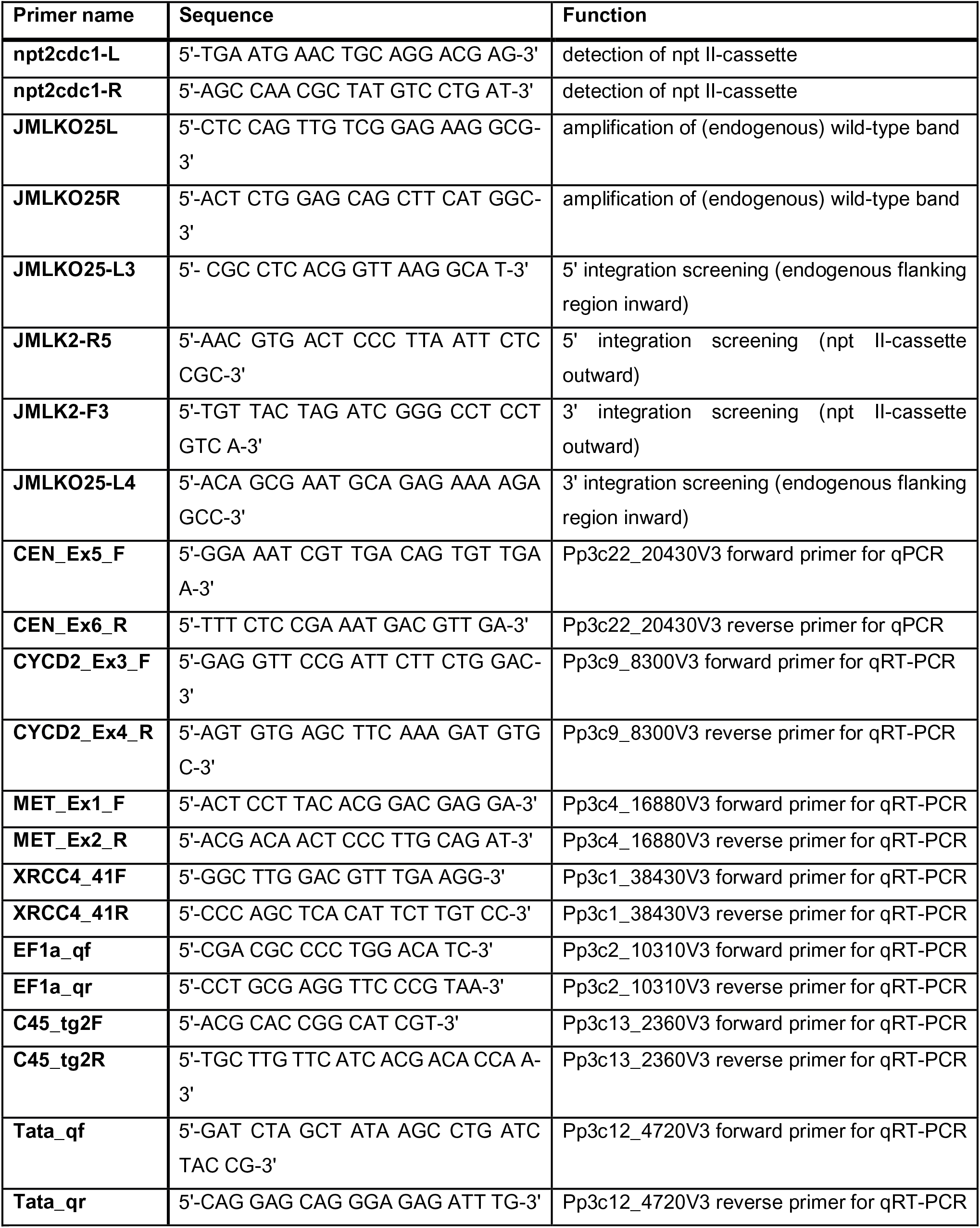
List of primers used for PCR screening of transformed Physcomitrella plants and for quantitative real-time PCR (qRT-PCR) of Physcomitrella cDNA.

**Supplementary Figure S2.**
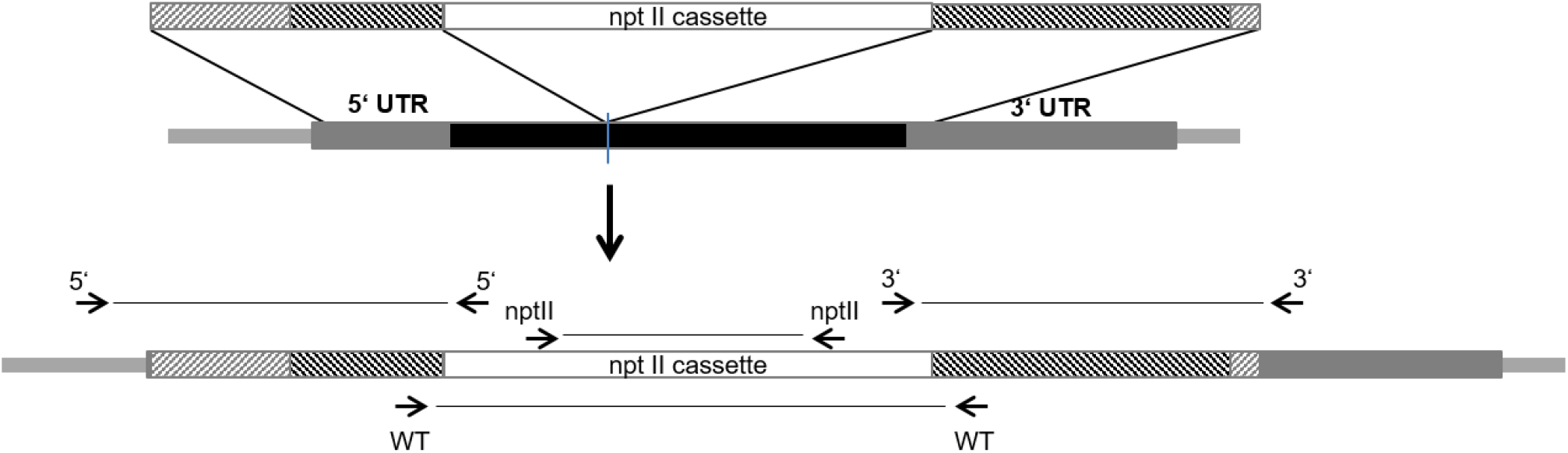
Schematic overview of knock-out construct pRKO25.2 integrated in the Physcomitrella genome. The location of primers used for PCR analysis of wild-type locus, 5’ integration and 3’ integration is shown.

**Supplementary Table T2.** Overview of the analyses performed with different Physcomitrella lines and sample sources. Differentially expressed genes were computed for different factors (time; ploidy at a certain time-point after protoplast isolation and transfection; ploidy) using protonema (PN), freshly isolated protoplasts (0h) as well as protoplasts at 4h and 24h after transfection.

| Factor | Lines | Comparisons |  |  |
| --- | --- | --- | --- | --- |
|  |  | First microarray | Second microarray | SuperSAGE |
| Time | WT | 1h vs. 0h<br>4h vs. 0h<br>6h vs. 0h<br>24h vs. 0h<br>72h vs. 0h |  | 4h vs. 0h<br>24h vs. 0h |
| Time | Haploid A |  | 4h vs. 0h<br>24h vs. 0h<br>PN vs. 0h | 4h vs. 0h<br>24h vs. 0h<br>PN vs. 0h |
| Time | Diploid A |  | 4h vs. 0h<br>24h vs. 0h<br>PN vs. P0h | 4h vs. 0h<br>24h vs. 0h<br>PN vs. 0h |
| Ploidy (time-point specific) | Diploid A vs. Haploid A |  | 0h<br>4h<br>24h<br>PN | 0h<br>4h<br>24h<br>PN |
| Ploidy (time-point specific) | Diploid A, Diploid B vs. WT, Haploid A |  | 0h<br>4h<br>24h<br>PN |  |
| Ploidy (time-point specific) | Diploid A vs. WT |  |  | 0h<br>4h<br>24h |
| Ploidy | Diploid A vs. WT, Haploid A |  |  | Two-factor analysis of: 0h, 4h, 24h and PN |
| Ploidy | Diploid A vs. WT |  |  | Two-factor analysis of: 0h, 4h and 24h |

**Supplementary Table T3.** Overview of SuperSAGE libraries used for analysis of differential gene expression. Libraries are constructed from 26 nt long reads as obtained by SuperSAGE technology and are mainly characterized by cell line and cell type of the material. Samples were taken from freshly isolated protoplasts (0h) and protoplasts at 4h and 24h after transfection. The libraries marked with * are mock transformants not exposed to foreign DNA. They were subjected to the whole transfection procedure but using water instead of the GT construct. The number of sequenced reads of the different libraries ranged from approximately 1.49 x 10^6^ to 24.21 x 10^6^ per library while the number of distinct sequences per library ranged from approximately 0.18 x 10^6^ to 1.46 x 10^6^. An average of 81.8% of the sequences was mapped to the V3.3 annotation of the Physcomitrella genome (Lang et al., 2018). On average 84.6% of the mapped reads resulted in a valid alignment as defined per featureCounts parameters (Supplementary Table T5). The majority of reads in the SuperSAGE data were highly abundant. More than half of the reads represented single sequences that were present more than 500 times on average per library while more than 10% of the reads belonged to sequences appearing over 10,000 times on average per library.

| Line | Ploidy | Tissue | Replicates |
| --- | --- | --- | --- |
| Diploid A | Diploid | protoplast 0h | 1 |
| Diploid A | Diploid | protoplast 4h | 1 |
| Diploid A | Diploid | protoplast 24h | 1 |
| Diploid A | Diploid | protonema | 2 |
| Haploid A | Haploid | protoplast 0h | 1 |
| Haploid A | Haploid | protoplast 4h | 1 |
| Haploid A | Haploid | protoplast 24h | 1 |
| Haploid A | Haploid | protonema | 2 |
| WT | Haploid | protoplast 0h | 1 |
| WT | Haploid | protoplast 4h | 1 |
| WT | Haploid | protoplast 4h | 2* |
| WT | Haploid | protoplast 24h | 1 |
| WT | Haploid | protoplast 24h | 2* |

**Supplementary Table T4.** Applied input parameters for the HISAT2 (v2.0.3, Kim et al., 2015) tool on the Galaxy platform (Freiburg Galaxy instance, http://galaxy.uni-freiburg.de, Afgan et al., 2016). The SuperSAGE reads were mapped to the V3.3 assembly of the Physcomitrella genome and known splice sites were provided in a gene annotation file in gff3 format.

| Input Parameter | Value |
| --- | --- |
| Input data format | Fastq |
| Single end or paired reads? | Single |
| Primary alignments | 50 |
| Alignment options | Defaults |
| Input options | Defaults |
| Scoring options: Function type | Constant |
| Scoring options: Constant term (y) | -2 |
| Scoring options: Coefficient (z) | 0 |
| Scoring options: Set match bonus | 2 |
| Scoring options: Maximum mismatch penalty | 6 |
| Scoring options: Minimum mismatch penalty | 2 |
| Scoring options: Ambiguous read penalty | 1 |
| Scoring options: Maximum soft-clipping penalty | 2 |
| Scoring options: Minimum soft-clipping penalty | 1 |
| Scoring options: Read gap open penalty | 5 |
| Scoring options: Read gap extend penalty | 3 |
| Scoring options: Reference gap open penalty | 5 |
| Scoring options: Reference gap extend penalty | 3 |
| Spliced alignment parameters: Penalty for canonical splice sites | 0 |
| Spliced alignment parameters: Penalty for non-canonical splice sites | 3 |
| Spliced alignment parameters: Penalty for long introns with canonical splice sites | Constant |
| Spliced alignment parameters: Constant term (y) | 0 |
| Spliced alignment parameters: Coefficient (z) | 0 |
| Spliced alignment parameters: Penalty for long introns with noncanonical splice sites | Constant |
| Spliced alignment parameters: Constant term (y) | -8 |
| Spliced alignment parameters: Coefficient (z) | 1 |
| Spliced alignment parameters :Minimum intron length | 20 |
| Spliced alignment parameters: Maximum intron length | 500000 |
| Spliced alignment parameters: Specify strand-specific information | Second Strand<br>(F/FR) |
| Disable spliced alignment | False |
| Transcriptome assembly reporting | Use default<br>reporting |
| Paired alignment parameters | Defaults |

**Supplementary Table T5.** Applied input parameters for the featureCounts tool (v1.4.6.p5, Liao et al., 2014) on the Galaxy platform (Freiburg Galaxy instance, http://galaxy.uni-freiburg.de, Afgan et al., 2016). Count tables were constructed from the output of the HISAT2 (v2.0.3) mapping tool using the V3.3 Physcomitrella annotation files.

| Input Parameter | Value |
| --- | --- |
| Output format | Gene-ID "\t" read-count |
| featureCounts parameters: GFF feature type filter | Exon |
| featureCounts parameters: GFF gene identifier | gene_id |
| featureCounts parameters: Allow read to contribute to multiple features | False |
| featureCounts parameters: Strand specific protocol | Unstranded |
| featureCounts parameters: Count multi-mapping reads/fragments | True |
| featureCounts parameters: Minimum read quality | 0 |
| Paired-end reads options: Count fragments instead of reads | True |
| Paired-end reads options: Check paired-end distance | False |
| Paired-end reads options: Minimum fragment/template length | 50 |
| Paired-end reads options: Maximum fragment/template length | 600 |
| Paired-end reads options: only allow fragments with both reads aligned | False |
| Paired-end reads options: Exclude chimeric fragments | True |
| On feature level | False |

**Supplementary Excel Sheet 1** Ploidy and time dependent differentially expressed genes (DEGs).

**Supplementary Excel Sheet 2** Enriched GO terms in haploid and diploid lines.

**Supplementary Table T6.** Number and overlap of genes being upregulated or downregulated between protonema samples or protoplasts at 4h or 24h after transfection versus freshly isolated protoplasts (0h). DEGs were computed from microarray (M) and SuperSAGE (S) data of two haploid lines (WT, Haploid A) and a diploid line (Diploid A). DEGs from the microarray experiment were determined with the Expressionist Analyst Pro software and were filtered for |log2 fold change| > 1 and p < 0.001. The SuperSAGE data analysis was performed with GFOLD and DEGs were filtered for a GFOLD(0.01) value of < -1 or > 1.

| Line | 4h versus 0h protoplast |  |  |  | 24h versus 0h protoplast |  |  |  | Protonema versus protoplast (0h) |  |  |  | Overlap<br>4h vs. 0h<br>with 24h<br>vs. 0h |
| --- | --- | --- | --- | --- | --- | --- | --- | --- | --- | --- | --- | --- | --- |
|  | total | up | down | down<br>[%] | total | up | down | down<br>[%] | total | up | down | down<br>[%] | total |
| (S) WT | 1148 | 444 | 704 | 61.32 | 4000 | 1167 | 2833 | 70.83 | - | - | - | - | 963 |
| (S) Haploid A | 3002 | 978 | 2024 | 67.42 | 3265 | 860 | 2405 | 73.66 | 740 | 74 | 666 | 90.00 | 1701 |
| (S) Diploid A | 1453 | 491 | 962 | 66.21 | 3949 | 2905 | 1044 | 26.44 | 545 | 3 | 542 | 99.45 | 1085 |
| (M) Haploid A | 262 | 48 | 214 | 81.67 | 2273 | 1365 | 908 | 39.95 | 1453 | 366 | 1087 | 74.81 | 162 |
| (M) Diploid A | 309 | 260 | 49 | 15.86 | 1894 | 1575 | 319 | 16.84 | 1674 | 958 | 716 | 42.77 | 223 |
| Combined (M,<br>S) Haploid A | 3142 | 995 | 2147 | 68.33 | 4663 | 1879 | 2784 | 59.70 | 1696 | 424 | 1272 | 75.00 | 1925 |
| Combined (M,<br>S) Haploid A<br>and WT | 3823 | 1297 | 2526 | 66.07 | 6698 | 2451 | 4247 | 63.41 | 1696 | 424 | 1272 | 75.00 | 2774 |
| Combined (M,<br>S) Diploid A | 1628 | 653 | 975 | 59.89 | 4922 | 3714 | 1208 | 24.54 | 1860 | 960 | 900 | 48.39 | 1246 |

**Supplementary Figure S3.**
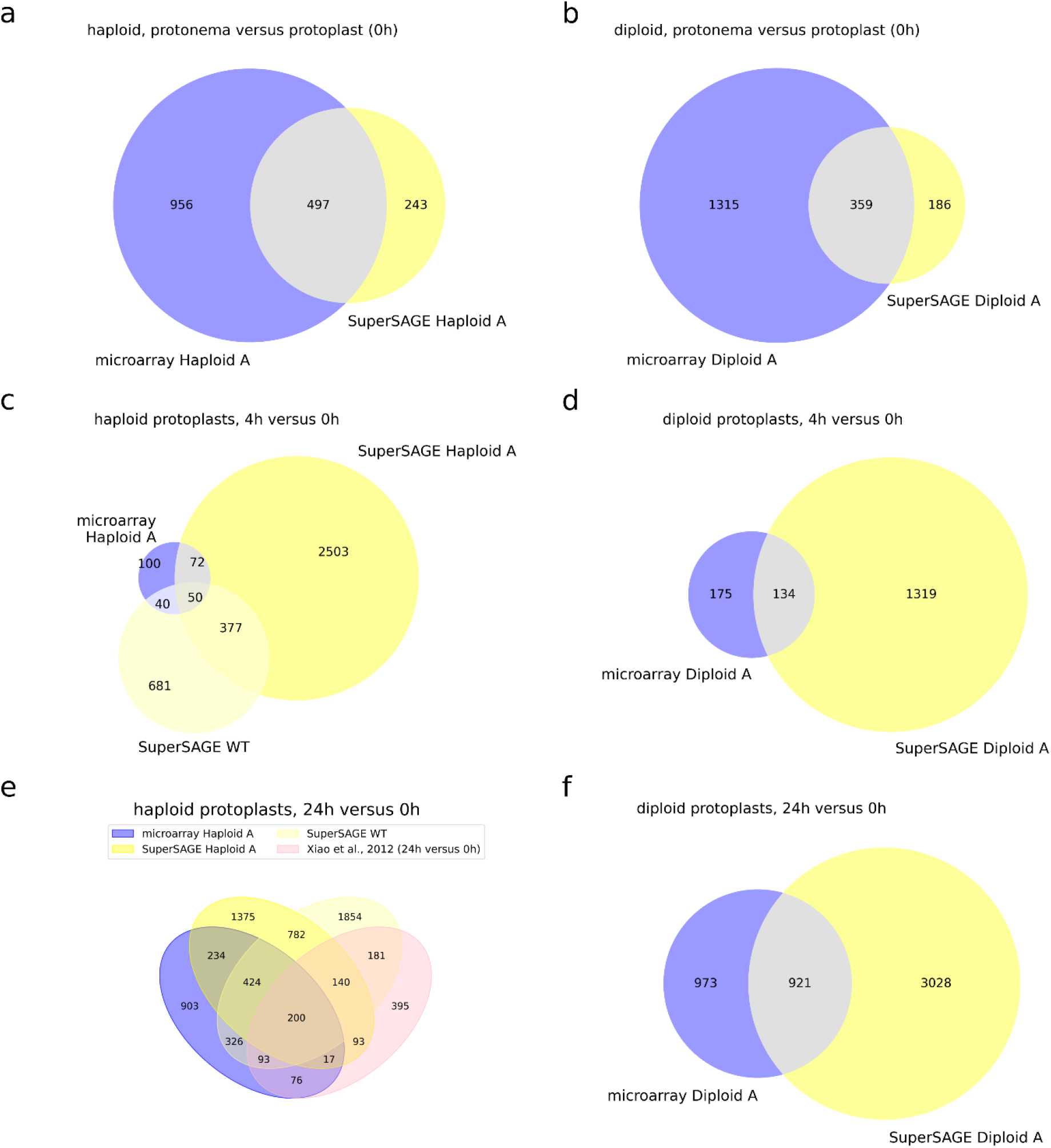
Overlap of differentially expressed genes (DEGs) from microarray and SuperSAGE data. DEGs were computed from protonema samples (a, b), protoplasts 4h after transfection (c, d) and protoplasts 24h after transfection (d, e), respectively, versus freshly isolated (0h) protoplasts using two haploid lines (a, c, d) and one diploid line (b, d, e). Results from microarray data are depicted in blue and results from SuperSAGE data in yellow. For our identified DEGs from the haploid lines at 24h (e) the overlap to DEGs from 24h old versus freshly isolated protoplasts identified by Xiao et al. (2012) is shown (pink).

**Supplementary Table T7.**
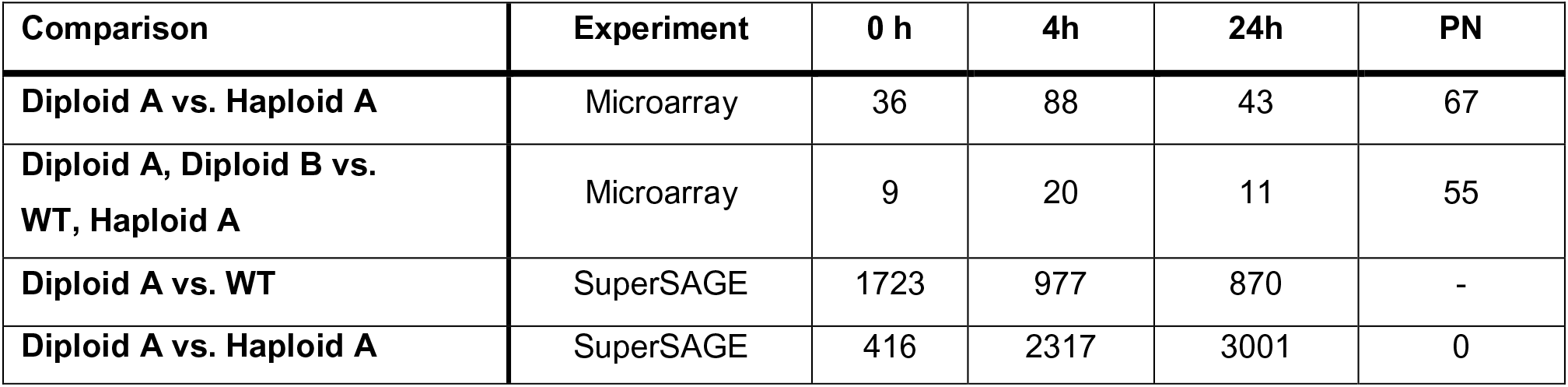
Number of differentially expressed genes (DEGs) between diploid and haploid lines in freshly isolated protoplasts (0h), in protoplasts at 4h and 24h after transfection as well as in protonema cells (PN). Comparisons were performed on microarray and SuperSAGE data from different lines. DEGs from the microarray experiment were determined with the Expressionist Analyst Pro software and filtered for a |log2 fold change| > 1 and p < 0.001. The SuperSAGE data analysis was performed with GFOLD and DEGs were filtered for a GFOLD(0.01) value of < -1 or > 1.

**Supplementary Figure S4.**
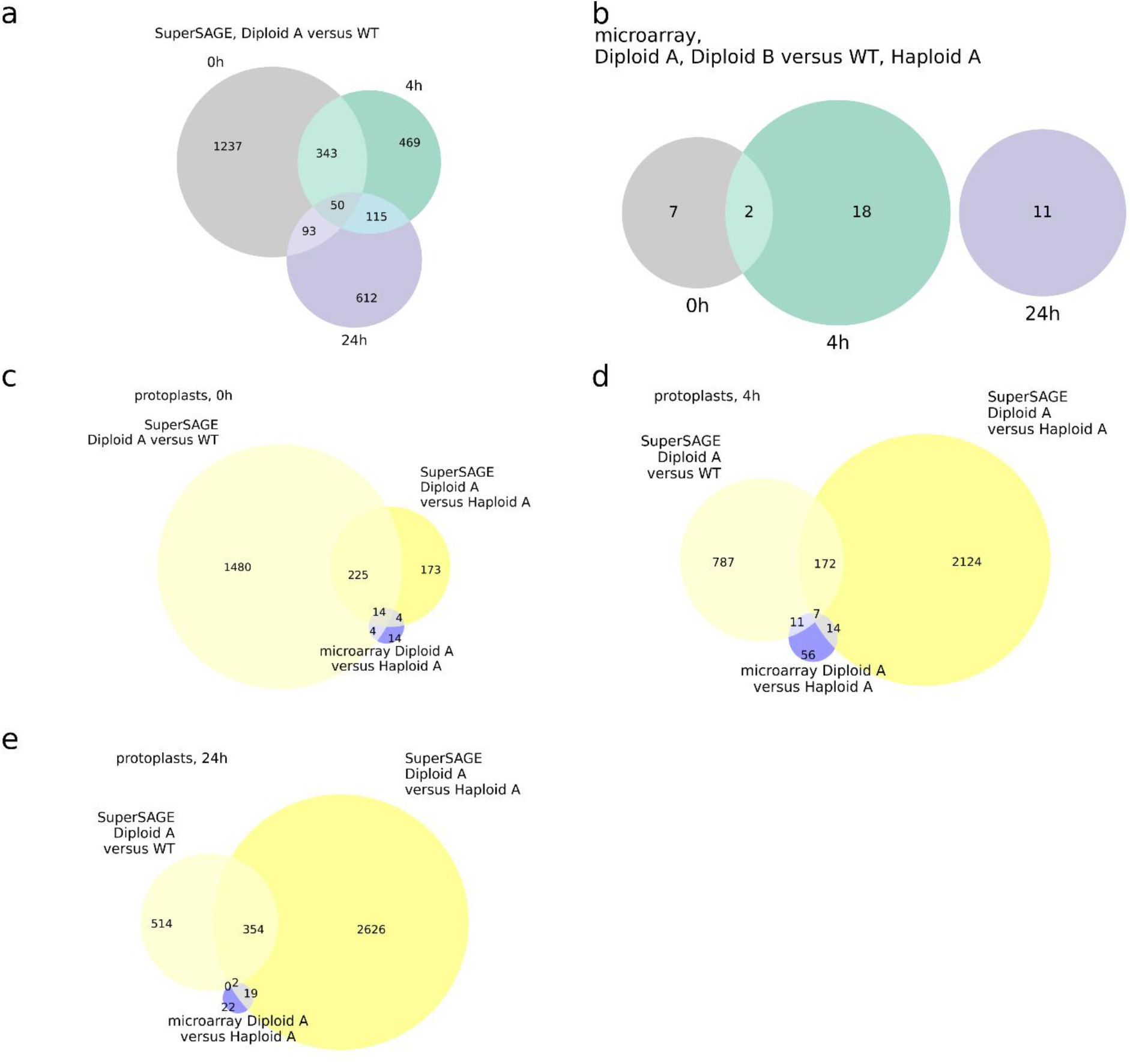
Overlap of differentially expressed genes (DEGs) identified from pairwise comparison between haploid and diploid protoplast cells. Shown is the overlap between DEGs at different protoplast ages (grey: freshly isolated protoplasts (0h), green: protoplasts at 4h after transfection, purple: protoplasts at 24h after transfection) from Diploid A versus WT using SuperSAGE libraries (a) and from Diploid A and Diploid B versus WT and Haploid A using microarray libraries (b). Further, the overlap of DEGs from microarray (blue) and SuperSAGE data (yellow) at 0h (c), 4h after transfection (d) and 24h after transfection (e) is illustrated. DEGs from the microarray experiment were determined with the Expressionist Analyst Pro software and were filtered for |log2 fold change| > 1 and p < 0.001. The SuperSAGE data analysis was performed with GFOLD and DEGs were filtered for a GFOLD(0.01) value of < -1 or > 1.

**Supplementary Figure S5.**
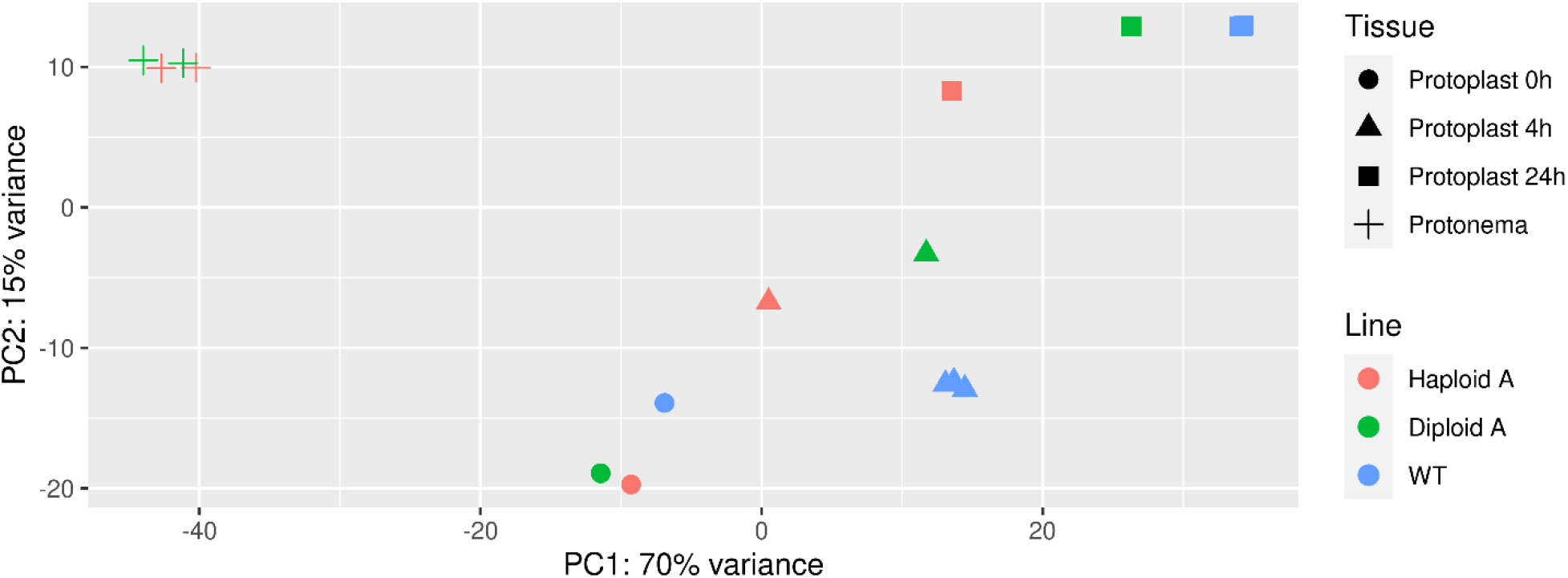
Comparison of variance between samples used for SuperSAGE library construction via principal component analysis (PCA). Similarity of haploid and diploid lines and tissue (including protoplast age) is shown. Tissue type and protoplast age are the main factors of the variance encountered.

**Supplementary Table T8.** Overview of the differentially expressed genes (DEGs) with the highest fold changes in gene expression between the diploid line Diploid A and the haploid WT. Listed are the 10 DEGs with the strongest upregulation as well as the 10 DEGs with the strongest downregulation. Statistical analysis was performed with DESeq2 using a two-factor design to test for ploidy-dependent expression factoring in the differences derived from different developmental stages. Only SuperSAGE libraries of protoplast samples were included in the analysis (Supplementary Table T3). Expression fold changes are given for diploid cells in comparison to haploid cells. The annotation is based on Phytozome (v12.1.5; Goodstein et al. 2012).

| V33 ID | Description | Log2 fold change | Adjusted p value |
| --- | --- | --- | --- |
| Pp3c23_940 | PF07470 - Glycosyl Hydrolase Family 88 (Glyco_hydro_88) (1 of 5) | 3.50 | 1.64E-15 |
| Pp3c22_10130 | K05605 - 3-hydroxyisobutyryl-CoA hydrolase (HIBCH) (1 of 4) | 3.44 | 2.66E-17 |
| Pp3c4_17790 | not annotated | 3.16 | 1.04E-17 |
| Pp3c4_25770 | PTHR28039:SF4 - CHALCONE--FLAVONONE ISOMERASE 3-RELATED (1 of 2) | 3.08 | 1.32E-14 |
| Pp3c2_21940 | not annotated | 3.02 | 1.65E-10 |
| Pp3c1_18940 | K10775 - phenylalanine ammonia-lyase (E4.3.1.24) (1 of 12) | 2.61 | 2.76E-11 |
| Pp3c4_24450 | not annotated | 2.55 | 2.54E-21 |
| Pp3c2_30370 | not annotated | 2.53 | 8.83E-10 |
| Pp3c25_14520 | not annotated | 2.52 | 1.57E-20 |
| Pp3c6_6560 | PTHR21495:SF39 - DIRIGENT PROTEIN 16-RELATED (1 of 8) | 2.51 | 6.04E-08 |
| Pp3c4_20120 | 1.1.3.9 - Galactose oxidase / Beta-galactose oxidase (1 of 36) | -2.17 | 2.56E-05 |
| Pp3c3_16280 | PTHR31867:SF30 - EXPANSIN-A5 (1 of 29) | -2.18 | 5.18E-05 |
| Pp3c24_12370 | PTHR11069 - GLUCOSYLCERAMIDASE (1 of 1) | -2.20 | 2.08E-06 |
| Pp3c21_14285 | not annotated | -2.24 | 1.21E-04 |
| Pp3c21_14239 | not annotated | -2.28 | 8.23E-05 |
| Pp3c12_15190 | PTHR31989:SF39 - NAC DOMAIN-CONTAINING PROTEIN 12 (1 of 10) | -2.31 | 2.50E-08 |
| Pp3c13_12570 | PTHR24067:SF41 - AKT-INTERACTING PROTEIN (1 of 1) | -2.39 | 1.14E-06 |
| Pp3c21_14233 | KOG0472 - Leucine-rich repeat protein (1 of 34) | -2.44 | 1.75E-05 |
| Pp3c18_19690 | PTHR31867:SF30 - EXPANSIN-A5 (1 of 29) | -2.63 | 1.51E-11 |
| Pp3c4_19000 | K02977 - small subunit ribosomal protein S27Ae (RP-S27Ae, RPS27A) (1 of 4) | -3.25 | 3.96E-44 |

**Supplementary Table T9.** Overview of the differentially expressed genes (DEGs) with the highest fold changes in gene expression between a diploid line (Diploid A) and two haploid lines (WT, Haploid A). Listed are the 10 DEGs with the strongest upregulation as well as the 10 DEGs with the strongest downregulation. Statistical analysis was performed with DESeq2 using a two-factor design to test for ploidy-dependent expression factoring in the differences derived from different developmental stages. All 17 SuperSAGE libraries of protoplast and protonema samples (Supplementary Table T3) were included in the analysis. Expression fold changes are given for diploid cells in comparison to haploid cells. The annotation is based on Phytozome (v12.1.5; Goodstein et al., 2012).

| V33 ID | Description | Log2 fold change | Adjusted p value |
| --- | --- | --- | --- |
| Pp3c20_15490 | PTHR11360 - MONOCARBOXYLATE TRANSPORTER (1 of 1) | 1.56 | 2.21E-04 |
| Pp3c4_17790 | not annotated | 1.53 | 2.20E-03 |
| Pp3c23_940 | PF07470 - Glycosyl Hydrolase Family 88 (Glyco_hydro_88) (1 of 5) | 1.52 | 2.90E-03 |
| Pp3c3_810 | PTHR11514//PTHR11514:SF57 - MYC // SUBFAMILY NOT NAMED (1 of 5) | 1.52 | 9.75E-06 |
| Pp3c13_18730 | not annotated | 1.51 | 9.51E-05 |
| Pp3c7_20720 | PF02743//PF13519 - Cache domain (Cache_1) // von Willebrand factor type A domain (VWA_2) (1 of 1) | 1.47 | 2.00E-03 |
| Pp3c4_20080 | K01087 - trehalose 6-phosphate phosphatase (otsB) (1 of 6) | 1.47 | 1.09E-04 |
| Pp3c11_2950 | K09753 - cinnamoyl-CoA reductase (CCR) (1 of 2) | 1.44 | 2.05E-03 |
| Pp3c15_4030 | K12640 - brassinosteroid-6-oxidase 2 (CYP85A2, BR6OX2) (1 of 1) | 1.44 | 2.47E-03 |
| Pp3c4_2660 | PF00847 - AP2 domain (AP2) (1 of 150) | 1.43 | 3.63E-05 |
| Pp3c4_30220 | PTHR19139:SF90 - AQUAPORIN PIP1-5-RELATED (1 of 3) | -1.53 | 2.43E-03 |
| Pp3c1_42230 | not annotated | -1.55 | 2.33E-03 |
| Pp3c1_42250 | not annotated | -1.55 | 2.33E-03 |
| Pp3c1_42420 | not annotated | -1.55 | 2.33E-03 |
| Pp3s350_10 | not annotated | -1.55 | 2.33E-03 |
| Pp3c2_8570 | not annotated | -1.58 | 2.29E-04 |
| Pp3c4_20120 | 1.1.3.9 - Galactose oxidase / Beta-galactose oxidase (1 of 36) | -1.69 | 2.37E-04 |
| Pp3c20_8180 | K14500 - BR-signalling kinase (BSK) (1 of 4) | -1.71 | 9.51E-05 |
| Pp3c17_19460 | PF04885 - Stigma-specific protein, Stig1 (Stig1) (1 of 7) | -1.85 | 4.87E-05 |
| Pp3c3_16280 | PTHR31867:SF30 - EXPANSIN-A5 (1 of 29) | -2.28 | 2.04E-08 |

**Supplementary Table T10.**
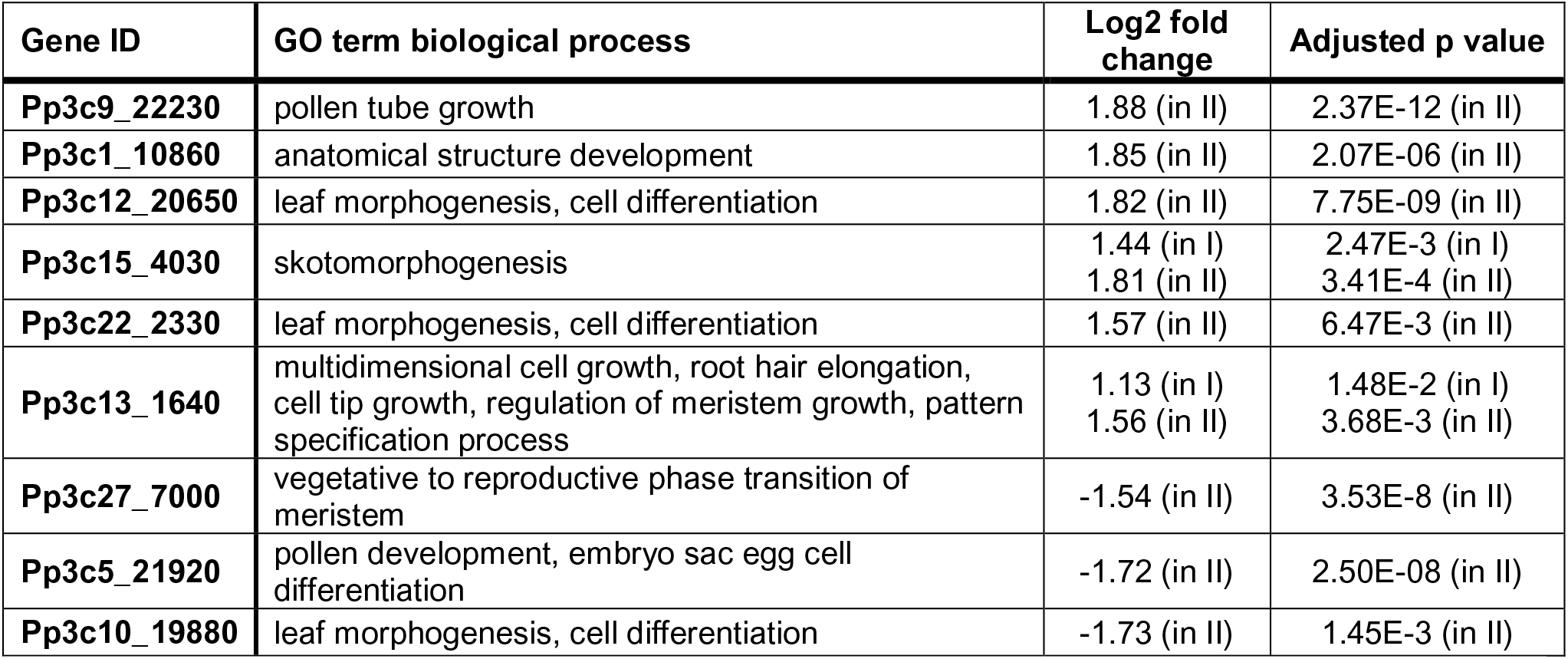
Differentially expressed genes (DEGs) between diploid and haploid cells involved in developmental or growth processes. From the two-factor analyses DEGs with a |log2 fold change| > 1.5 were selected if they had a biological process GO term assigned considering development process (GO:0032502), growth (GO:0040007) or any of their child terms. Only these GO terms are listed in the table. DEGs in the first two-factor analysis (I) were computed from Diploid A versus WT and Haploid A using protoplast and protonema samples whereas DEGs in the second two-factor analysis (II) were computed from Diploid A versus WT using only protoplast samples.

| Gene ID | GO term biological process | Log2 fold change | Adjusted p value |
| --- | --- | --- | --- |
| **Pp3c9_22230** | pollen tube growth | 1.88 (in II) | 2.37E-12 (in II) |
| **Pp3c1_10860** | anatomical structure development | 1.85 (in II) | 2.07E-06 (in II) |
| **Pp3c12_20650** | leaf morphogenesis, cell differentiation | 1.82 (in II) | 7.75E-09 (in II) |
| **Pp3c15_4030** | skotomorphogenesis | 1.44 (in I)<br>1.81 (in II) | 2.47E-3 (in I)<br>3.41E-4 (in II) |
| **Pp3c22_2330** | leaf morphogenesis, cell differentiation | 1.57 (in II) | 6.47E-3 (in II) |
| **Pp3c13_1640** | multidimensional cell growth, root hair elongation, cell tip growth, regulation of meristem growth, pattern specification process | 1.13 (in I)<br>1.56 (in II) | 1.48E-2 (in I)<br>3.68E-3 (in II) |
| **Pp3c27_7000** | vegetative to reproductive phase transition of meristem | -1.54 (in II) | 3.53E-8 (in II) |
| **Pp3c5_21920** | pollen development, embryo sac egg cell differentiation | -1.72 (in II) | 2.50E-08 (in II) |
| **Pp3c10_19880** | leaf morphogenesis, cell differentiation | -1.73 (in II) | 1.45E-3 (in II) |

**Supplementary Figure S6.**
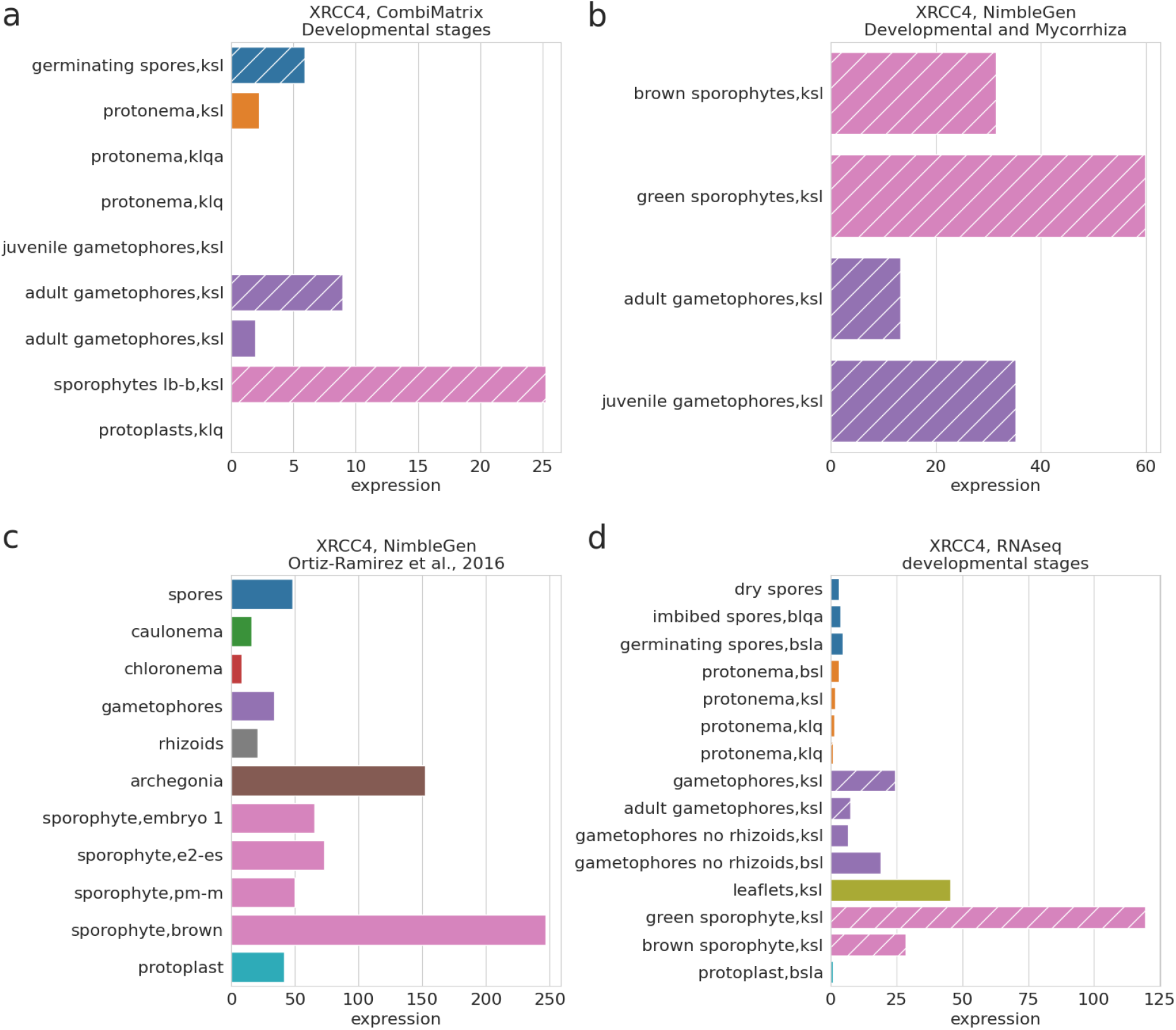
XRCC4 expression in the Reute and Gransden Physcomitrella ecotypes at different developmental stages. Expression values were obtained from the PEATmoss website and only the four datasets containing values for the gene expression in the sporophyte were selected. Gene expression in the datasets CombiMatrix Developmental stages (a), NimbleGen Developmental and Mycorrhiza (b), NimbleGen Ortiz-Ramírez et al., 2016 (c) and RNAseq developmental stages (d) is shown. In the *NimbleGen Developmental and Mycorrhiza* dataset mycorrhiza exudate and heat treated samples were not considered. Hatched bars represent expression values from the Reute ecotype. Abbreviations: BlqA = BCDA (ammonium) liquid, Bsl = BCD solid, BslA = BCDA (ammonium) solid, Klq = Knop liquid, KlqA = Knop liquid ammonium, Ksl = Knop solid. Colours of source material: Spores in blue, protonema in orange, gametophores in purple, sporophyte in pink, protoplast in turquoise, archegonia in brown, rhizoids in grey, chloronema in red, caulonema in green, leaflets in olive.

**Supplementary Figure S7.**
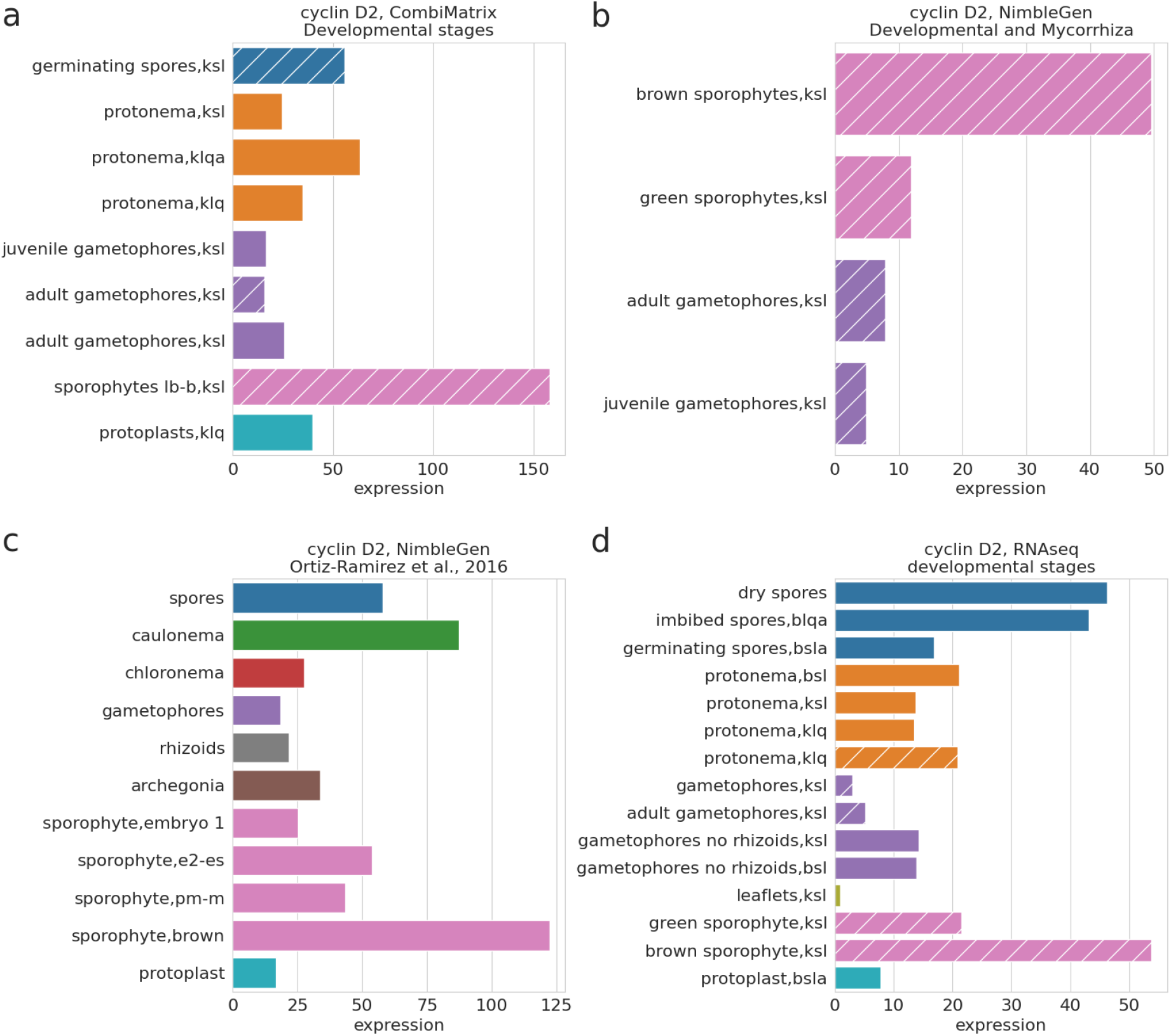
Cyclin D2 expression in the Reute and Gransden Physcomitrella ecotypes at different developmental stages. Expression values were obtained from the PEATmoss website and only the four datasets containing values for the gene expression in the sporophyte were selected. The gene expression in the datasets CombiMatrix Developmental stages (a), NimbleGen Developmental and Mycorrhiza (b), NimbleGen Ortiz-Ramírez et al., 2016 (c) and RNAseq developmental stages (d) is shown. In the *NimbleGen Developmental and Mycorrhiza* dataset mycorrhiza exudate and heat treated samples were not considered. Hatched bars represent expression values from the Reute ecotype. Abbreviations: BlqA = BCDA (ammonium) liquid, Bsl = BCD solid, BslA = BCDA (ammonium) solid, Klq = Knop liquid, KlqA = Knop liquid ammonium, Ksl = Knop solid. Colours of source material: Spores in blue, protonema in orange, gametophores in purple, sporophyte in pink, protoplast in turquoise, archegonia in brown, rhizoids in grey, chloronema in red, caulonema in green, leaflets in olive.

**Supplementary Figure S8.**
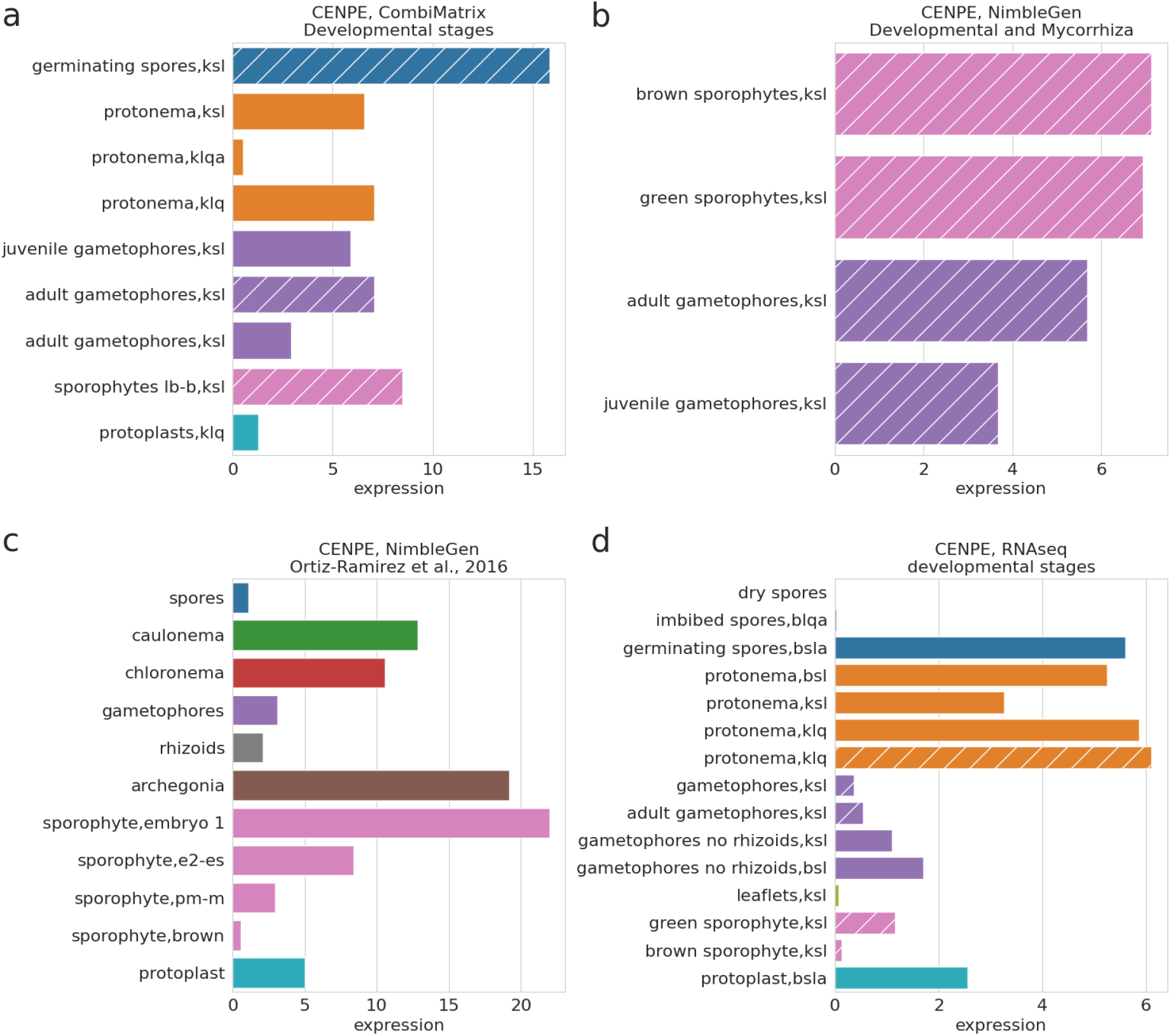
CENPE expression in the Reute and Gransden Physcomitrella ecotypes at different developmental stages. Expression values were obtained from the PEATmoss website and only the four datasets containing values for the gene expression in the sporophyte were selected. The gene expression in the datasets CombiMatrix Developmental stages (a), NimbleGen Developmental and Mycorrhiza (b), NimbleGen Ortiz-Ramírez et al., 2016 (c) and RNAseq developmental stages (d) is shown. In the *NimbleGen Developmental and Mycorrhiza* dataset mycorrhiza exudate and heat treated samples were not considered. Hatched bars represent expression values from the Reute ecotype Abbreviations: BlqA = BCDA (ammonium) liquid, Bsl = BCD solid, BslA = BCDA (ammonium) solid, Klq = Knop liquid, KlqA = Knop liquid ammonium, Ksl = Knop solid. Colours of source material: Spores in blue, protonema in orange, gametophores in purple, sporophyte in pink, protoplast in turquoise, archegonia in brown, rhizoids in grey, chloronema in red, caulonema in green, leaflets in olive.

**Supplementary Figure S9.**
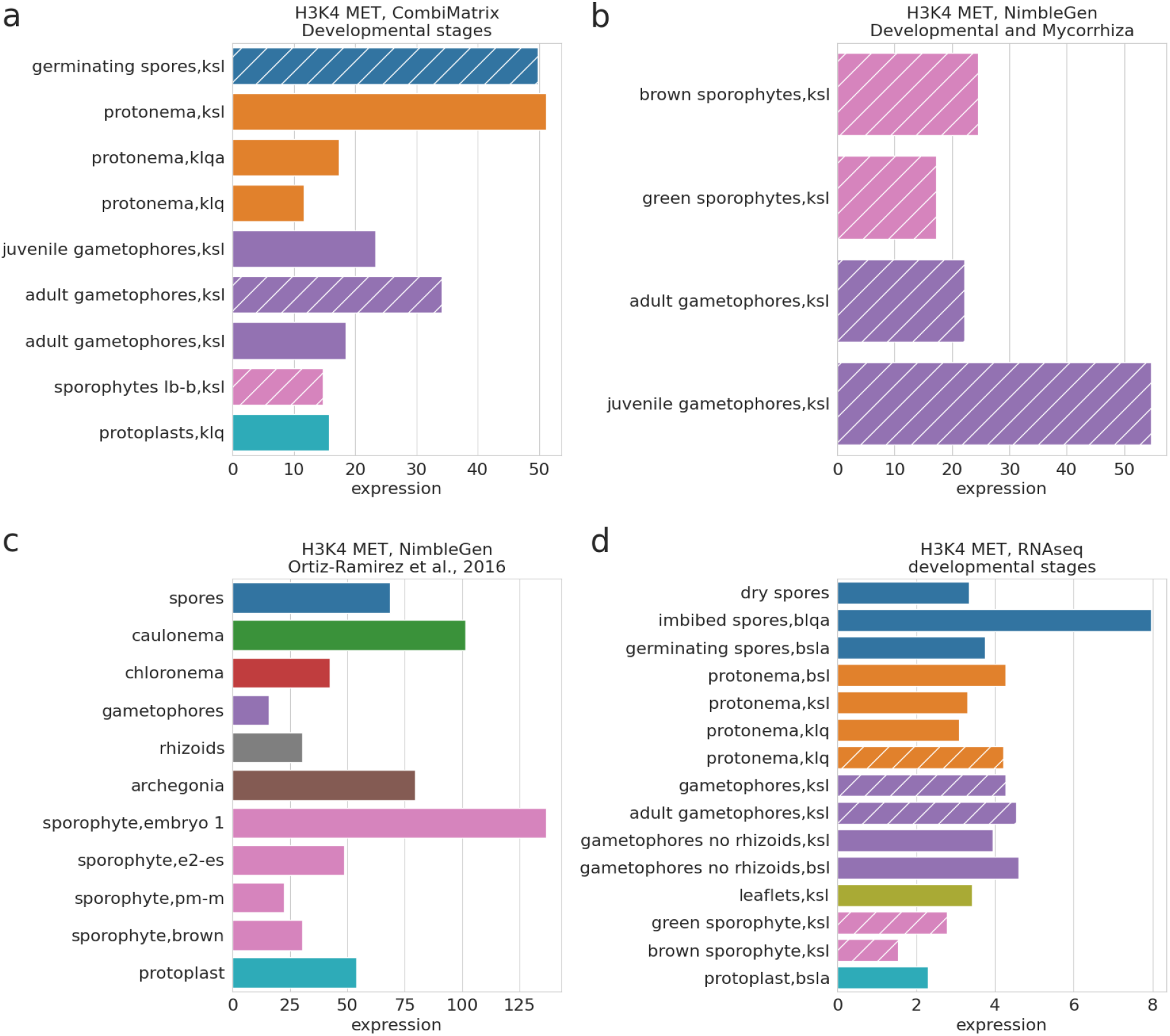
H3K4 expression in the Reute and Gransden Physcomitrella ecotypes at different developmental stages. Expression values were obtained from the PEATmoss website and only the four datasets containing values for the gene expression in the sporophyte were selected. The gene expression in the datasets CombiMatrix Developmental stages (a), NimbleGen Developmental and Mycorrhiza (b), NimbleGen Ortiz-Ramírez et al., 2016 (c) and RNAseq developmental stages (d) is shown. In the *NimbleGen Developmental and Mycorrhiza* dataset mycorrhiza exudate and heat treated samples were not considered. Hatched bars represent expression values from the Reute ecotype Abbreviations: BlqA = BCDA (ammonium) liquid, Bsl = BCD solid, BslA = BCDA (ammonium) solid, Klq = Knop liquid, KlqA = Knop liquid ammonium, Ksl = Knop solid. Colours of source material: Spores in blue, protonema in orange, gametophores in purple, sporophyte in pink, protoplast in turquoise, archegonia in brown, rhizoids in grey, chloronema in red, caulonema in green, leaflets in olive.

## References

Abel WO, Knebel W, Koop H-U, Marienfeld JR, Quader H, Reski R, Schnepf E, Spörlein B (1989) A cytokinin-sensitive mutant of the moss, *Physcomitrella patens*, defective in chloroplast division. Protoplasma 152:1–13. https://doi.org/10.1007/BF01354234

Abrieu A, Kahana JA, Wood KW, Cleveland DW (2000) CENP-E as an essential component of the mitotic checkpoint *in vitro*. Cell 102:817–826. https://doi.org/10.1016/S0092-8674(00)00070-2

Adams KL, Wendel JF (2005) Polyploidy and genome evolution in plants. Curr Opin Plant Biol 8:135–141. https://doi.org/10.1016/j.pbi.2005.01.001

Afgan E, Baker D, van den Beek M, Blankenberg D, Bouvier D, Čech M, Chilton J, Clements D, Coraor N, Eberhard C, et al. (2016) The Galaxy platform for accessible, reproducible and collaborative biomedical analyses: 2016 update. Nucleic Acids Res 44, W3–W10. https://doi.org/10.1093/nar/gkw343

Amano K, Sago H, Uchikawa C, Suzuki T, Kotliarova SE, Nukina N, Epstein CJ, Yamakawa K (2004) Dosage-dependent over-expression of genes in the trisomic region of Ts1Cje mouse model for Down syndrome. Hum Mol Gen 13:1333–1340. https://doi.org/10.1093/hmg/ddh154

Andalis AA, Storchova Z, Styles C, Galitski T, Pellman D, Fink GR (2004) Defects arising from whole-genome duplications in *Saccharomyces cerevisiae*. Genetics 167:1109–1121. https://doi.org/10.1534/genetics.104.029256

Andrews S (2010) FastQC: A quality control tool for high throughput sequence data. Available online at: http://www.bioinformaticsbabrahamacuk/projects/fastqc/

Attaran E, Major IT, Cruz JA, Rosa BA, Koo AJK, Chen J, Kramer DM, He SY, Howe GA (2014) Temporal dynamics of growth and photosynthesis suppression in response to jasmonate signaling. Plant Physiol 165:1302–1314. https://doi.org/10.1104/pp.114.239004

Bakr A, Oing C, Köcher S, Borgmann K, Dornreiter I, Petersen C, Dikomey E, Mansour WY (2015) Involvement of ATM in homologous recombination after end resection and RAD51 nucleofilament formation. Nucleic Acids Res 43:3154–3166. https://doi.org/10.1093/nar/gkv160

Baldi P, Long AD (2001) A Bayesian framework for the analysis of microarray expression data: Regularized t-test and statistical inferences of gene changes. Bioinformatics 17:509–519. https://doi.org/10.1093/bioinformatics/17.6.509

Barone P, Wu E, Lenderts B, Anand A, Gordon-Kamm W, Svitashev S, Kumar S (2020) Efficient gene targeting in maize using inducible CRISPR-Cas9 and marker-free donor template. Mol Plant 13:1219–1227. https://doi.org/10.1016/j.molp.2020.06.008

Beetham PR, Kipp PB, Sawycky XL, Arntzen CJ, May GD (1999) A tool for functional plant genomics: chimeric RNA/DNA oligonucleotides cause *in vivo* gene-specific mutations. Proc Natl Acad Sci USA 96:8774–8778. https://doi.org/10.1073/pnas.96.15.8774

Behling AH, Shepherd LD, Cox MP (2020) The importance and prevalence of allopolyploidy in Aotearoa New Zealand. J R Soc NZ 50:189–210. https://doi.org/10.1080/03036758.2019.1676797

Beike AK, von Stackelberg M, Schallenberg-Rüdinger M, Hanke ST, Follo M, Quandt D, McDaniel SF, Reski R, Tan BC, Rensing SA (2014) Molecular evidence for convergent evolution and allopolyploid speciation within the *Physcomitrium-Physcomitrella* species complex. BMC Evol Biol 14:158. https://doi.org/10.1186/1471-2148-14-158

Beike AK, Lang D, Zimmer AD, Wüst F, Trautmann D, Wiedemann G, Beyer P, Decker EL, Reski R (2015) Insights from the cold transcriptome of *Physcomitrella patens*: global specialization pattern of conserved transcriptional regulators and identification of orphan genes involved in cold acclimation. New Phytol 205:869–881. https://doi.org/10.1111/nph.13004

Ben-David U, Amon A (2020) Context is everything: aneuploidy in cancer. Nat Rev Genet 21:44–62. https://doi.org/10.1038/s41576-019-0171-x

Birchler JA, Veitia RA (2012) Gene balance hypothesis: connecting issues of dosage sensitivity across biological disciplines. Proc Natl Acad Sci USA 109:14746–14753. https://doi.org/10.1073/pnas.1207726109

Britt AB, May GD (2003) Re-engineering plant gene targeting. Trends Plant Sci 8:90–95. https://doi.org/10.1016/S1360-1385(03)00002-5

Brouwer I, Sitters G, Candelli A, Heerema SJ, Heller I, de Melo AJ, Zhang H, Normanno D, Modesti M, Peterman EJG, et al. (2016) Sliding sleeves of XRCC4-XLF bridge DNA and connect fragments of broken DNA. Nature 535:566–569. https://doi.org/10.1038/nature18643

Chang HHY, Pannunzio NR, Adachi N, Lieber MR (2017) Non-homologous DNA end joining and alternative pathways to double-strand break repair. Nat Rev Mol Cell Biol 18:495–506. https://doi.org/10.1038/nrm.2017.48

Cheng F, Wu J, Cai X, Liang J, Freeling M, Wang X (2018) Gene retention, fractionation and subgenome differences in polyploid plants. Nat Plants 4:258–268. https://doi.org/10.1038/s41477-018-0136-7

Cohen H, Fait A, Tel-Zur N (2013) Morphological, cytological and metabolic consequences of autopolyploidization in *Hylocereus* (Cactaceae) species. BMC Plant Biol 13:173. https://doi.org/10.1186/1471-2229-13-173

Cohen SP, Leach JE (2019) Abiotic and biotic stresses induce a core transcriptome response in rice. Sci Rep 9:6273. https://doi.org/10.1038/s41598-019-42731-8

Collonnier C, Guyon-Debast A, Maclot F, Mara K, Charlot F, Nogué F (2017) Towards mastering CRISPR-induced gene knock-in in plants: Survey of key features and focus on the model *Physcomitrella patens*. Methods 121–122:103–117. https://doi.org/10.1016/j.ymeth.2017.04.024

Comai L (2005) The advantages and disadvantages of being polyploid. Nat Rev Genet 6:836–846. https://doi.org/10.1038/nrg1711

Davies G, Henrissat B (1995) Structures and mechanisms of glycosyl hydrolases. Structure 3:853–859. https://doi.org/10.1016/S0969-2126(01)00220-9

Davis AJ, Chen DJ (2013) DNA double strand break repair via non-homologous end-joining. Transl Cancer Res 2:130–143. https://doi.org/10.3978/j.issn.2218-676X.2013.04.02

Decker EL, Wiedemann G, Reski R (2015) Gene Targeting for Precision Glyco-Engineering: Production of Biopharmaceuticals Devoid of Plant-Typical Glycosylation in Moss Bioreactors. In: Castilho A. (eds) Glyco-Engineering. Methods in Molecular Biology, vol 1321. Humana Press, New York, NY. https://doi.org/10.1007/978-1-4939-2760-9_15

del Pozo JC, Ramirez-Parra E (2015) Whole genome duplications in plants: an overview from *Arabidopsis*. J Exp Bot 66:6991–7003. https://doi.org/10.1093/jxb/erv432

Dong C, Beetham P, Vincent K, Sharp P (2006) Oligonucleotide-directed gene repair in wheat using a transient plasmid gene repair assay system. Plant Cell Rep 25:457–465.

Doyle JJ, Flagel LE, Paterson AH, Rapp RA, Soltis DE, Soltis PS, Wendel JF (2008) Evolutionary genetics of genome merger and doubling in plants. Annu Rev Genet 42:443–461. https://doi.org/10.1146/annurev.genet.42.110807.091524

Duijf PHG, Schultz N, Benezra R (2013) Cancer cells preferentially lose small chromosomes. Int J Cancer 132:2316–2326. https://doi.org/10.1002/ijc.27924

Egener T, Granado J, Guitton MC, Hohe A, Holtorf H, Lucht JM, Rensing SA, Schlink K, Schulte J, Schween G, et al. (2002) High frequency of phenotypic deviations in *Physcomitrella patens* plants transformed with a gene-disruption library. BMC Plant Biol 2:6. https://doi.org/10.1186/1471-2229-2-6

El Kelish A, Zhao F, Heller W, Durner J, Winkler JB, Behrendt H, Traidl-Hoffmann C, Horres R, Pfeifer M, Frank U, et al. (2014) Ragweed (*Ambrosia artemisiifolia*) pollen allergenicity: SuperSAGE transcriptomic analysis upon elevated CO_2_ and drought stress. BMC Plant Biol 14:176. https://doi.org/10.1186/1471-2229-14-176

Feng J, Meyer CA, Wang Q, Liu JS, Shirley Liu X, Zhang Y (2012) GFOLD: a generalized fold change for ranking differentially expressed genes from RNA-seq data. Bioinformatics 28:2782–2788. https://doi.org/10.1093/bioinformatics/bts515

Fernandez-Pozo N, Haas FB, Meyberg R, Ullrich KK, Hiss M, Perroud PF, Hanke S, Kratz V, Powell AF, Vesty EF, et al. (2020), PEATmoss (*Physcomitrella* Expression Atlas Tool): a unified gene expression atlas for the model plant *Physcomitrella patens*. Plant J 102:165–177. https://doi.org/10.1111/tpj.14607

Frank W, Decker EL, Reski R (2005) Molecular tools to study *Physcomitrella patens*. Plant Biol 7:220–227. https://doi.org/10.1055/s-2005-865645

Gautier L (2010) An intuitive Python interface for Bioconductor libraries demonstrates the utility of language translators. BMC Bioinformatics 11:S11. https://doi.org/10.1186/1471-2105-11-S12-S11

Girke T, Schmidt H, Zähringer U, Reski R, Heinz E (1998) Identification of a novel delta 6-acyl-group desaturase by targeted gene disruption in *Physcomitrella patens*. Plant J 15:39–48. https://doi.org/10.1046/j.1365-313X.1998.00178.x

Goffová I, Vágnerová R, Peška V, Franek M, Havlová K, Holá M, Zachová D, Fojtová M, Cuming A, Kamisugi Y, et al. (2019) Roles of RAD51 and RTEL1 in telomere and rDNA stability in *Physcomitrella patens*. Plant J 98:1090–1105. https://doi.org/10.1046/j.1365-313x.1998.00178.x

Goodstein DM, Shu S, Howson R, Neupane R, Hayes RD, Fazo J, Mitros T, Dirks W, Hellsten U, Putnam N, et al. (2012) Phytozome: a comparative platform for green plant genomics. Nucleic Acids Res 40:D1178–D1186. https://doi.org/10.1093/nar/gkr944

Graham TGW, Walter JC, Loparo JJ (2016) Two-stage synapsis of DNA ends during non-homologous end joining. Mol Cell 61:850–858. https://doi.org/10.1016/j.molcel.2016.02.010

Gu C, Guo ZH, Hao PP, Wang GM, Jin ZM, Zhang SL (2017) Multiple regulatory roles of AP2/ERF transcription factor in angiosperm. Bot Stud 58:6. https://doi.org/10.1186/s40529-016-0159-1

Gu N, Tamada Y, Imai A, Palfalvi G, Kabeya Y, Shigenobu S, Ishikawa M, Angelis KJ, Chen C, Hasebe M (2020) DNA damage triggers reprogramming of differentiated cells into stem cells in Physcomitrella. Nat Plants 6:1098–1105. https://doi.org/10.1038/s41477-020-0745-9

Gurrieri L, Fermani S, Zaffagnini M, Sparla F, Trost P (2021) Calvin-Benson cycle regulation is getting complex. Trends Plant Sci. https://doi.org/10.1016/j.tplants.2021.03.008

Guo M, Davis D, Birchler JA (1996) Dosage effects on gene expression in a maize ploidy series. Genetics 142:1349–1355.

Guyon-Debast A, Rossetti P, Charlot F, Epert A, Neuhaus JM, Schaefer DG, Nogué F (2019) The XPF-ERCC1 complex is essential for genome stability and is involved in the mechanism of gene targeting in *Physcomitrella patens*. Front Plant Sci 10:588. https://doi.org/10.3389/fpls.2019.00588

Harris CR, Millman KJ, van der Walt SJ, Gommers R, Virtanen P, Cournapeau D, Wieser E, Taylor J, Berg S, Smith NJ, et al. (2020) Array programming with NumPy. Nature 585:357–362. https://doi.org/10.1038/s41586-020-2649-2

Hayashi K, Horie K, Hiwatashi Y, Kawaide H, Yamaguchi S, Hanada A, Nakashima T, Nakajima M, Mander LN, Yamane H, et al. (2010) Endogenous diterpenes derived from *ent*-kaurene, a common gibberellin precursor, regulate protonema differentiation of the moss *Physcomitrella patens*. Plant Physiol 153:1085–1097. https://doi.org/10.1104/pp.110.157909

He P, Shan L, Sheen J (2007) The use of protoplasts to study innate immune responses. Methods Mol Bio 354:1–9. https://doi.org/10.1385/1-59259-966-4:1

Heck MA, Lüth VM, van Gessel N, Krebs M, Kohl M, Prager A, Joosten H, Decker EL, Reski R (2021) Axenic *in vitro* cultivation of 19 peat moss (*Sphagnum* L) species as a resource for basic biology, biotechnology, and paludiculture. New Phytol 229:861–876. https://doi.org/10.1111/nph.16922

Hellemans J, Mortier G, De Paepe A, Speleman F, Vandesompele J (2007) qBase relative quantification framework and software for management and automated analysis of real-time quantitative PCR data. Genome Biol 8:R19. https://doi.org/10.1186/gb-2007-8-2-r19

Heyer WD, Ehmsen KT, Liu J (2010) Regulation of homologous recombination in eukaryotes. Annu Rev Genet 44:113–139. https://doi.org/10.1146/annurev-genet-051710-150955

Hiss M, Laule O, Meskauskiene RM, Arif MA, Decker EL, Erxleben A, Frank W, Hanke ST, Lang D, Martin A, et al. (2014) Large-scale gene expression profiling data for the model moss *Physcomitrella patens* aid understanding of developmental progression, culture and stress conditions. Plant J 79:530–539. https://doi.org/10.1111/tpj.12572

Hohe A, Egener T, Lucht JM, Holtorf H, Reinhard C, Schween G, Reski R (2004) An improved and highly standardised transformation procedure allows efficient production of single and multiple targeted gene-knockouts in a moss, *Physcomitrella patens*. Curr Genet 44:339–347. https://doi.org/10.1007/s00294-003-0458-4

Horst NA, Katz A, Pereman I, Decker EL, Ohad N, Reski R (2016) A single homeobox gene triggers phase transition, embryogenesis and asexual reproduction. Nat Plants 2:15209. https://doi.org/10.1038/nplants.2015.209

Horst NA, Reski R (2016) Alternation of generations – unravelling the underlying molecular mechanism of a 165-year-old botanical observation. Plant Biol 18:549–551. https://doi.org/10.1111/plb.12468

Hunter JD (2007) Matplotlib: a 2D graphics environment. Comp Sci Eng 9:90–95.

Hyun MW, Yun YH, Kim JY, Kim SH (2011) Fungal and plant phenylalanine ammonia-lyase. Mycobiology 39:257–265. https://doi.org/10.5941/MYCO.2011.39.4.257

Iiizumi S, Kurosawa A, So S, Ishii Y, Chikaraishi Y, Ishii A, Koyama H, Adachi N (2008) integration: implications for gene targeting. Nucleic Acids Res 36:6333–6342. https://doi.org/10.1093/nar/gkn649

Ishizaki K, Johzuka-Hisatomi Y, Ishida S, Iida S, Kohchi T (2013) Homologous recombination-mediated gene targeting in the liverwort *Marchantia polymorpha* L. Sci Rep 3:1532. https://doi.org/10.1038/srep01532

Kamisugi Y, Cuming AC, Cove DJ (2005) Parameters determining the efficiency of gene targeting in the moss *Physcomitrella patens*. Nucleic Acids Res 33:e173. https://doi.org/10.1093/nar/gni172

Kamisugi Y, Schlink K, Rensing SA, Schween G, von Stackelberg M, Cuming AC, Reski R, Cove DJ (2006) The mechanism of gene targeting in *Physcomitrella patens*: homologous recombination, concatenation and multiple integration. Nucleic Acids Res 34:6205–6214. https://doi.org/10.1093/nar/gkl832

Kamisugi Y, Schaefer DG, Kozak J, Charlot F, Vrielynck N, Holá M, Angelis KJ, Cuming AC, Nogué F (2012) MRE11 and RAD50, but not NBS1, are essential for gene targeting in the moss *Physcomitrella patens*. Nucleic Acids Res 40:3496–3510. https://doi.org/10.1093/nar/gkr1272

Kamisugi Y, Whitaker JW, Cuming AC (2016) The transcriptional response to DNA-double-strand breaks in *Physcomitrella patens*. PLoS One 11:e0161204. https://doi.org/10.1371/journal.pone.0161204

Kim D, Langmead B, Salzberg SL (2015) HISAT: a fast spliced aligner with low memory requirements. Nat Methods 12:357–360. https://doi.org/10.1038/nmeth.3317

Lang D, Ullrich KK, Murat F, Fuchs J, Jenkins J, Haas FB, Piednoel M, Gundlach H, Van Bel M, Meyberg R, et al. (2018) The *Physcomitrella patens* chromosome-scale assembly reveals moss genome structure and evolution. Plant J 93:515–533. https://doi.org/10.1111/tpj.13801

Lee JW, Blanco L, Zhou T, Garcia-Diaz M, Bebenek K, Kunkel TA, Wang Z, Povirk LF (2004) Implication of DNA polymerase λ in alignment-based gap filling for nonhomologous DNA end joining in human nuclear extracts. J Biol Chem 279:805–811. https://doi.org/10.1074/jbc.M307913200

Letourneau A, Santoni FA, Bonilla X, Sailani MR, Gonzalez D, Kind J, Chevalier C, Thurman R, Sandstrom RS, Hibaoui Y, et al. (2014) Domains of genome-wide gene 47 expression dysregulation in Down’s syndrome. Nature 508:345–350. https://doi.org/10.1038/nature13200

Li H, Handsaker B, Wysoker A, Fennell T, Ruan J, Homer N, Marth G, Abecasis G, Durbin R, 1000 Genome Project Data Processing Subgroup (2009) The Sequence Alignment/Map format and SAMtools. Bioinformatics 25:2078–2079. https://doi.org/10.1093/bioinformatics/btp352

Li XC, Tye BK (2011) Ploidy dictates repair pathway choice under DNA replication stress. Genetics 187:1031–1040. https://doi.org/10.1534/genetics.110.125450

Liao Y, Smyth GK, Shi W (2014) featureCounts: an efficient general purpose program for assigning sequence reads to genomic features. Bioinformatics 30:923–930. https://doi.org/10.1093/bioinformatics/btt656

Liu B, Wendel JF (2003) Epigenetic phenomena and the evolution of plant allopolyploids. Mol Phylogenet Evol 29:365–379. https://doi.org/10.1016/s1055-7903(03)00213-6

Love MI, Huber W, Anders S (2014) Moderated estimation of fold change and dispersion for RNA-seq data with DESeq2. Genome Biol 15:550. https://doi.org/10.1186/s13059-014-0550-8

Manghwar H, Lindsey K, Zhang X, Jin S, (2019) CRISPR/Cas system: recent advances and future prospects for genome editing. Trends Plant Sci 24:1102–1125. https://doi.org/10.1016/j.tplants.2019.09.006

Mao Y, Botella JR, Liu Y, Zhu JK (2019) Gene editing in plants: progress and challenges. Natl Sci Rev 6:421–437. https://doi.org/10.1093/nsr/nwz005

Mara K, Charlot F, Guyon-Debast A, Schaefer DG, Collonnier C, Grelon M, Nogué F (2019) POLQ plays a key role in the repair of CRISPR/Cas9-induced double-stranded breaks in the moss *Physcomitrella patens*. New Phytol 222:1380–1391. https://doi.org/10.1111/nph.15680

Maréchal A, Zou L (2013) DNA damage sensing by the ATM and ATR kinases. Cold Spring Harb Perspect Biol 5:a012716. https://doi.org/10.1101/cshperspect.a012716

Markmann-Mulisch U, Wendeler E, Zobell O, Schween G, Steinbiss HH, Reiss B (2007) Differential requirements for RAD51 in *Physcomitrella patens* and *Arabidopsis thaliana* development and DNA damage repair. Plant Cell 19:3080–3089.

Marowa P, Ding A, Kong Y (2016) Expansins: roles in plant growth and potential applications in crop improvement. Plant Cell Rep 35:949–965. https://doi.org/10.1007/s00299-016-1948-4

Martens M, Horres R, Wendeler E, Reiss B (2020) The Importance of ATM and ATR in *Physcomitrella* patens DNA damage repair, development, and gene targeting. Genes 11:752. https://doi.org/10.3390/genes11070752

Martin A, Lang D, Hanke ST, Mueller SJX, Sarnighausen E, Vervliet-Scheebaum M, Reski (2009) Targeted gene knockouts reveal overlapping functions of the five *Physcomitrella patens* FtsZ isoforms in chloroplast division, chloroplast shaping, cell patterning, plant development, and gravity sensing. Molecular Plant 2:1359–1372. https://doi.org/10.1093/mp/ssp076

Matsumura H, Yoshida K, Luo S, Kimura E, Fujibe T, Albertyn Z, Barrero RA, Krüger DH, Kahl G, Schroth GP, et al. (2010) High-Throughput SuperSAGE for digital gene expression analysis of multiple samples using next generation sequencing. PloS One 5:e12010. https://doi.org/10.1371/journal.pone.0012010

Matsuoka Y (2011) Evolution of polyploid *Triticum* wheats under cultivation: the role of domestication, natural hybridization and allopolyploid speciation in their diversification Plant Cell Physiol 52:750–764. https://doi.org/10.1093/pcp/pcr018

McKinney W (2010) Data structures for statistical computing in python. In Proc of the 9th Python in Science Conf 445:56-61.

Medina R, Johnson M, Liu Y, Wilding N, Hedderson TA, Wickett N, Goffinet B (2018) Evolutionary dynamism in bryophytes: phylogenomic inferences confirm rapid radiation in the moss family Funariaceae. Mol Phylogenet Evol 120:240–247. https://doi.org/10.1016/j.ympev.2017.12.002

Medina R, Johnson MG, Liu Y, Wickett NJ, Shaw AJ, Goffinet B (2019) Phylogenomic delineation of *Physcomitrium* (Bryophyta: Funariaceae) based on targeted sequencing of nuclear exons and their flanking regions rejects the retention of *Physcomitrella*, *Physcomitridium* and *Aphanorrhegma*. J Syst Evol 57:404–417. https://doi.org/10.1111/jse.12516

Mengiste T, Paszkowski J (1999) Prospects for the precise engineering of plant genomes by homologous recombination. Biol Chem 380:749–758. https://doi.org/10.1515/BC.1999.095

Okuzaki A, Toriyama K (2004) Chimeric RNA/DNA oligonucleotide-directed gene targeting in rice. Plant Cell Rep 22:509–512. https://doi.org/10.1007/s00299-003-0698-2

Ortiz-Ramírez C, Hernandez-Coronado M, Thamm A, Catarino B, Wang M, Dolan L, Feijó JA, Becker JD (2016) A transcriptome atlas of *Physcomitrella patens* provides insights into the evolution and development of land plants. Mol Plant 9:205–220. https://doi.org/10.1016/j.molp.2015.12.002

Ostendorf AK, van Gessel N, Malkowsky Y, Sabovljevic MS, Rensing SA, Roth-Nebelsick A, Reski R (2021) Polyploidization within the Funariaceae – a key principle behind speciation, sporophyte reduction and the high variance of spore diameters? Bryophyte Div Evol, in press. https://doi.org/10.11646/bde.00.0.0

Otto SP (2007) The evolutionary consequences of polyploidy. Cell 131:452–62. https://doi.org/10.1016/j.cell.2007.10.022

Passerini V, Ozeri-Galai E, de Pagter MS, Donnelly N, Schmalbrock S, Kloosterman WP, Kerem B, Storchová Z (2016) The presence of extra chromosomes leads to genomic instability. Nat Comm 7:10754. https://doi.org/10.1038/ncomms10754

Puchta H (2002) Gene replacement by homologous recombination in plants. Plant Mol Biol 48:173–182. https://doi.org/10.1023/A:1013761821763

Qi X, Wu H Jiang H, Zhu J, Huang C, Zhang X, Liu C, Cheng B (2020) Conversion of a normal maize hybrid into a waxy version using *in vivo* CRISPR/Cas9 targeted mutation activity. Crop J 8:440–448. https://doi.org/10.1016/j.cj.2020.01.006

R core team (2020) R: A language and environment for statistical computing. R Found Statist Comput, Vienna, Austria, https://www.R-projectorg/

Rao X, Dixon RA (2017) Brassinosteroid mediated cell wall remodeling in grasses under abiotic stress. Front Plant Sci 8:806. https://doi.org/10.3389/fpls.2017.00806

Renny-Byfield S, Wendel JF (2014) Doubling down on genomes: polyploidy and crop plants. Am J Bot 101:1711–1725. https://doi.org/10.3732/ajb.1400119

Rensing SA, Lang D, Zimmer AD, Terry A, Salamov A, Shapiro H, Nishiyama T, Perroud PF, Lindquist EA, Kamisugi Y, et al. (2008) The *Physcomitrella* genome reveals evolutionary insights into the conquest of land by plants. Science 319:64–69. https://doi.org/10.1126/science.1150646

Resemann HC, Herrfurth C, Feussner K, Hornung E, Ostendorf AK, Gömann J, Mittag J, van Gessel N, de Vries J, Ludwig-Müller J, et al. (2021) Convergence of sphingolipid desaturation across over 500 million years of plant evolution. Nat Plants 7:219–232. https://doi.org/10.1038/s41477-020-00844-3

Reski R (1998a) Development, genetics and molecular biology of mosses. Bot Acta 111:1–15. https://doi.org/10.1111/j.1438-8677.1998.tb00670.x

Reski R (1998b) *Physcomitrella* and *Arabidopsis*: the David and Goliath of reverse genetics. Trends Plant Sci 3:209–210. https://doi.org/10.1016/S1360-1385(98)01257-6

Reski R, Abel WO (1985) Induction of budding on chloronemata and caulonemata of the moss, *Physcomitrella patens*, using isopentenyladenine. Planta 165:354–358. https://doi.org/10.1007/BF00392232

Richardt S, Timmerhaus G, Lang D, Qudeimat E, Corrêa LGG, Reski R, Rensing SA, Frank W (2010) Microarray analysis of the moss *Physcomitrella patens* reveals evolutionarily conserved transcriptional regulation of salt stress and abscisic acid signalling. Plant Mol Biol 72:27–45. https://doi.org/10.1007/s11103-009-9550-6

Samanta A, Das G, Das SK, (2011) Roles of flavonoids in plants. Int J Pharm Sci Technol 6:12–35.

Sattler MC, Carvalho CR, Clarindo WR (2016) The polyploidy and its key role in plant breeding. Planta 243:281–296. https://doi.org/10.1007/s00425-015-2450-x

Schaefer DG, Zrÿd JP (1997) Efficient gene targeting in the moss *Physcomitrella patens*. Plant J 11:1195–1206. https://doi.org/10.1046/j.1365-313x.1997.11061195.x

Schaefer DG, Delacote F, Charlot F, Vrielynck N, Guyon-Debast A, Le Guin S, Neuhaus JM, Doutriaux MP, Nogué F (2010) RAD51 loss of function abolishes gene targeting and de-represses illegitimate integration in the moss *Physcomitrella patens*. DNA Repair 9:526–533. https://doi.org/10.1016/j.dnarep.2010.02.001

Schneider M, Knuesting J, Birkholz O, Heinisch JJ, Scheibe R (2018) Cytosolic GAPDH as a redox-dependent regulator of energy metabolism. BMC Plant Biol 18:184. https://doi.org/10.1186/s12870-018-1390-6

Schindele A, Dorn A, Puchta H (2020) CRISPR/Cas brings plant biology and breeding into the fast lane. Curr Opin Biotech 61:7–14. https://doi.org/10.1016/j.copbio.2019.08.006

Schipper O, Schaefer D, Reski R, Fleming A (2002) Expansins in the bryophyte *Physcomitrella patens*. Plant Mol Biol 50:789–802. https://doi.org/10.1023/A:1019907207433

Schulte J, Erxleben A, Schween G, Reski R (2006) High throughput metabolic screen of *Physcomitrella* transformants. Bryologist 109:247–256. https://doi.org/10.1639/0007-2745(2006)109[247:HTMSOP]2.0.CO;2

Schween G, Fleig S, Reski R (2002) High-throughput-PCR screen of 15,000 transgenic *Physcomitrella* plants. Plant Mol Biol Rep 20:43–47. https://doi.org/10.1007/BF02801931

Schween G, Gorr G, Hohe A, Reski R (2003) Unique tissue-specific cell cycle in *Physcomitrella*. Plant Biol 5:50–58. https://doi.org/10.1055/s-2003-37984

Schween G, Egener T, Fritzowsky D, Granado J, Guitton MC, Hartmann N, Hohe A, Holtorf H, Lang D, Lucht JM, et al. (2005a) Large-scale analysis of 73 329 *Physcomitrella* plants transformed with different gene disruption libraries: production parameters and mutant phenotypes. Plant Biol 7:228–237. https://doi.org/10.1055/s-2005-837692

Schween G, Schulte J, Reski R, Hohe A (2005b) Effect of ploidy level on growth, differentiation, and morphology in *Physcomitrella patens*. Bryologist 108:27–35. https://doi.org/10.1639/0007-2745(2005)108[27:EOPLOG]2.0.CO;2

Sharma R, Tan F, Jung KH, Sharma MK, Peng Z, Ronald PC (2011) Transcriptional dynamics during cell wall removal and regeneration reveals key genes involved in cell wall development in rice. Plant Mol Biol 77:391–406. https://doi.org/10.1007/s11103-011-9819-4

Sharma S, Hicks JK, Chute CL, Brennan JR, Ahn JY, Glover TW, Canman CE (2012) REV1 and polymerase ζ facilitate homologous recombination repair. Nucleic Acids Res 40:682–691. https://doi.org/10.1093/nar/gkr769

Shi X, Zhang C, Ko DK, Chen ZJ (2015) Genome-wide dosage-dependent and - independent regulation contributes to gene expression and evolutionary novelty in plant polyploids. Mol Biol Evol 32:2351–2366. https://doi.org/10.1093/molbev/msv116

Smirnoff N, Arnaud D (2019) Hydrogen peroxide metabolism and functions in plants. New Phytol 221:1197–1214. https://doi.org/10.1111/nph.15488

Soltis PS, Marchant DB, Van de Peer Y, Soltis DE (2015) Polyploidy and genome evolution in plants. Curr Opin Genet Dev 35:119–125. https://doi.org/10.1016/j.pbi.2005.01.001

Soltis PS, Soltis DE (2016) Ancient WGD events as drivers of key innovations in angiosperms. Curr Opin Plant Biol 30:159–165. https://doi.org/10.1016/j.pbi.2016.03.015

Spoelhof JP, Soltis PS, Soltis DE (2017) Pure polyploidy: closing the gaps in autopolyploid research. J Syst Evol 55:340–352. https://doi.org/10.1111/jse.12253

Steinert J, Schiml S, Puchta H (2016) Homology-based double-strand break-induced genome engineering in plants. Plant Cell Rep 35:1429–1438. https://doi.org/10.1007/s00299-016-1981-3

Strepp R, Scholz S, Kruse S, Speth V, Reski R (1998) Plant nuclear gene knockout reveals a role in plastid division for the homolog of the bacterial cell division protein FtsZ, an ancestral tubulin. Proc Natl Acad Sci USA 95:4368–4373. https://doi.org/10.1073/pnas.95.8.4368

Strotbek C, Krinninger S, Frank W (2013) The moss *Physcomitrella patens*: methods and tools from cultivation to targeted analysis of gene function. Int J Dev Biol 57:553–564. https://doi.org/10.1387/ijdb.130189wf

The pandas development team (2020) pandas-dev/pandas: Pandas 1.0.3. Zenodo. https://doi.org/10.5281/zenodo.3715232

Torres EM, Williams BR, Amon A (2008) Aneuploidy: cells losing their balance. Genetics 179:737–746. https://doi.org/10.1534/genetics.108.090878

Trouiller B, Charlot F, Choinard S, Schaefer DG, Nogué F (2007) Comparison of gene targeting efficiencies in two mosses suggests that it is a conserved feature of bryophyte transformation. Biotechnol Lett 29:1591–1598. https://doi.org/10.1007/s10529-007-9423-5

van de Peer Y, Mizrachi E, Marchal K (2017) The evolutionary significance of polyploidy. Nat Rev Genet 18:411–424. https://doi.org/10.1038/nrg.2017.26

Van Rossum G, Drake FL (2009) Python 3 Reference Manual Scotts Valley, CA: CreateSpace

Vandenbussche F, Fierro AC, Wiedemann G, Reski R, Van Der Straeten D (2007) Evolutionary conservation of plant gibberellin signalling pathway components. BMC Plant Biol 7:65. https://doi.org/10.1186/1471-2229-7-65

Walden N, German DA, Wolf EM, Kiefer M, Rigault P, Huang XC, Kiefer C, Schmickl R, Franzke A, Neuffer B, et al. (2020) Nested whole-genome duplications coincide with diversification and high morphological disparity in Brassicaceae. Nat Comm 11:3795. https://doi.org/10.1038/s41467-020-17605-7

Waltz E (2016) Gene-edited CRISPR mushroom escapes US regulation. Nature 532:293. https://doi.org/10.1038/nature.2016.19754

Waskom M, and the seaborn development team (2020) mwaskom/seaborn: v0.10.1. Zenodo. https://doi.org/10.5281/zenodo.3767070

Watanabe K, Pacher M, Dukowic S, Schubert V, Puchta H, Schubert I (2009) The STRUCTURAL MAINTENANCE OF CHROMOSOMES 5/6 complex promotes sister chromatid alignment and homologous recombination after DNA damage *in Arabidopsis thaliana*. Plant Cell 21:2688–2699. https://doi.org/10.1105/tpc.108.060525

Weimer AK, Biedermann S, Harashima H, Roodbarkelari F, Takahashi N, Foreman J, Guan Y, Pochon G, Heese M, Van Damme D, et al. (2016) The plant-specific CDKB1-CYCB1 complex mediates homologous recombination repair in *Arabidopsis*. EMBO J 35:2068–2086. https://doi.org/10.15252/embj.201593083

Weiner J (2015) tagcloud: Tag Clouds R package version 06 https://CRANR-projectorg/package=tagcloud

Weterings E, Chen DJ (2008) The endless tale of non-homologous end-joining. Cell Res 18:114–124. https://doi.org/10.1038/cr.2008.3

Wiedemann G, van Gessel N, Köchl F, Hunn L, Schulze K, Maloukh L, Nogué F, Decker EL, Hartung F, Reski R (2018) RecQ helicases function in development, DNA repair, and gene targeting in *Physcomitrella patens*. Plant Cell 30:717–736. https://doi.org/10.1105/tpc.17.00632

Wolf L, Rizzini L, Stracke R, Ulm R, Rensing SA (2010) The molecular and physiological responses of *Physcomitrella patens* to ultraviolet-B radiation. Plant Physiol 153:1123– 1134. https://doi.org/10.1104/pp.110.154658

Wolffe AP, Matzke MA (1999) Epigenetics: regulation through repression. Science 286:481–486. https://doi.org/10.1126/science.286.5439.481

Wood AJ, Reski R, Frank W (2004) Isolation and characterization of ALDHIIA5, a novel non-phosphorylating GAPDH cDNA from *Physcomitrella patens*. Bryologist 107:385–387. https://doi.org/10.1639/0007-2745(2004)107[0385:IACOAA]2.0.CO;2

Wu JH, Ferguson AR, Murray BG, Jia Y, Datson PM, Zhang J (2012) Induced polyploidy dramatically increases the size and alters the shape of fruit in *Actinidia chinensis*. Ann Bot 109:169–179. https://doi.org/10.1093/aob/mcr256

Xiao L, Zhang L, Yang G, Zhu H, He Y (2012) Transcriptome of protoplasts reprogrammed into stem cells in *Physcomitrella patens*. PLoS One 7:e35961. https://doi.org/10.1371/journal.pone.0035961

Xu J, Wang X, Guo W (2015) The cytochrome P450 superfamily: key players in plant development and defense. J Integr Agric 14:1673–1686. https://doi.org/10.1016/S2095-3119(14)60980-1

Xu YP, Zhao Y, Song XY, Ye YF, Wang RG, Wang ZL, Ren XL, Cai XZ (2019) Ubiquitin extension protein UEP1 modulates cell death and resistance to various pathogens in tobacco. Phytopathology 109:1257–1269. https://doi.org/10.1094/PHYTO-06-18-0212-R

Yang X, Tu L, Zhu L, Fu L, Min L, Zhang X (2008) Expression profile analysis of genes involved in cell wall regeneration during protoplast culture in cotton by suppression subtractive hybridization and microarray. J Exp Bot 59:3661–3674. https://doi.org/10.1093/jxb/ern214

Yong B, Wang X, Xu P, Zheng H, Fei X, Hong Z, Ma Q, Miao Y, Yuan X, Jiang Y, et al. (2017) Isolation and abiotic stress resistance analyses of a catalase gene from *Ipomoea batatas* (L.) Lam. BioMed Res Int 2017:6847532. https://doi.org/10.1155/2017/6847532

Yu G, Wang LG, Han Y, He QY (2012) clusterProfiler: an R package for comparing biological themes among gene clusters. OMICS 16:284–287. https://doi.org/10.1089/omi.2011.0118

Yu P, He X, Baer M, Beirinckx S, Tian T, Moya YAT, Zhang X, Deichmann M, Frey FP Bresgen V, et al. (2021) Plant flavones enrich rhizosphere Oxalobacteraceae to improve maize performance under nitrogen deprivation. Nat Plants 7:481-499. https://doi.org/10.1038/s41477-021-00897-y

Zha S, Guo C, Boboila C, Oksenych V, Cheng H L, Zhang Y, Wesemann DR, Yuen G, Patel H, Goff PH, et al. (2011) ATM damage response and XLF repair factor are functionally redundant in joining DNA breaks. Nature 469:250–254. https://doi.org/10.1038/nature09604

Zhu T, Peterson DJ, Tagliani L, St Clair G, Baszczynski CL, Bowen B (1999) Targeted manipulation of maize genes *in vivo* using chimeric RNA/DNA oligonucleotides. Proc Natl Acad Sci USA 96:8768–8773. https://doi.org/10.1073/pnas.96.15.8768

